# Beyond clines: lineages and haplotype blocks in hybrid zones

**DOI:** 10.1101/043190

**Authors:** Alisa Sedghifar, Yaniv Brandvain, Peter Ralph

## Abstract

Hybrid zones formed between recently diverged populations offer an opportunity to study the mechanisms underlying reproductive isolation and the process of speciation. Here, we use a combination of analytical theory and explicit forward simulations to describe how selection against hybrid genotypes impacts patterns of introgression across genomic and geographic space. By describing how lineages move across the hybrid zone, in a model without coalescence, we add to modern understanding of how clines form and how parental haplotypes are broken up during introgression. Working with lineages makes it easy to see that clines form in about 1/*s* generations, where s is the strength of selection against hybrids, and linked clines persist over a genomic scale of 1/*T*, where *T* is the age, in generations, of the hybrid zone. Locally disadvantageous alleles tend to exist as small families, whose lineages trace back to the side from which they originated at speed 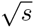 dispersal distances per generation. The lengths of continuous tracts of ancestry provide an additional source of information: blocks of ancestry surrounding incompatibilities can be substantially longer than the genome-wide average block length at the same spatial location, an observation that might be used to identify candidate targets of selection.

## Introduction

The process of speciation commonly involves populations diverging with relatively little gene flow (Coyne & Orr 2004). However, when formerly isolated populations come into contact before reproductive isolation is complete, some gene flow is possible. Interbreeding and migration between such populations creates a gradient of alleles derived from the two populations across geographic space, centered on their point of contact (reviewed by Barton & Hewitt 1985). If the populations are sufficiently diverged, this process leaves a distinct pattern of variation across the genome, in which long tracts of divergent haplotypes from each ancestral population are broken up by historical recombination events, forming chromosomal junctions in ancestry (Fisher 1954; Chapman & Thompson 2002; Baird *et al* 2003). These patterns of correlated ancestry in admixed populations have been previously used to infer histories of hybridization and admixture (e.g., Gravel 2012; Hellenthal *et al* 2014; Harris & Nielsen 2013; Sedghifar *et al* 2015). Here we show how natural selection against hybrid incompatibilities shapes patterns of coancestry around such loci.

Our work here builds on the large body of theory describing allele frequency clines in tension zones, and aims to describe transient dynamics with a focus on haplotype patterns. To do this, we take the reverse-time perspective, understanding the temporal dynamics of the system by describing how lineages tracing back from the hybrid zone moveme across space, conditioning on the frequencies of the selected allele and following them as they recombine between selected backgrounds (similar to Hudson & Kaplan 1988). Using a combination of theory and simulated hybrid zones, and extending the framework of Sedghifar *et al* (2015), we describe genome-wide patterns of coancestry as they relate to hybrid zone age, genetic distance from selected locus and geographic distance from hybrid zone center.

In particular, we examine how heterogeneous patterns of the ancestry length distribution across the genome may help identify putative targets of selection in hybridizing populations, providing another source of information in addition to gradients in allele frequencies used in current approaches (e.g., Porter *et al* 1997; Gompert *et al* 2012). Specifically, we find that selection against hybrid incompatibility loci changes the extent of correlated ancestry — that is, because selection rapidly removes incompatible alleles from heterospecific genomes, hybrid incompatibilities will be surrounded by disproportionately long ancestry blocks.

If hybrid ancestry at a given locus is disfavored, migrant haplotypes containing the selected allele will be removed rapidly from the population, preventing introgression of surrounding genomic regions. We therefore expect a deficit of short blocks of foreign ancestry surrounding the selected locus, with this effect becoming more pronounced further away from the center of a hybrid zone. As a corollary to this, conditional on having the ancestry that is at lower frequency (that is, being on the ‘wrong’ side of the hybrid zone), the length of unbroken ancestry surrounding the selected locus is expected to be relatively long when far away from the zone center. This is because an unfit haplotype is more likely to have been recently inherited from the other side of the hybrid zone, and therefore will not have as many ancestors of the locally common type as do neutral haplotypes. This reasoning is applicable to instances of both tension zones generated by intrinsic genetic incompatibilities, and ecotone models of extrinsic ecological selection (e.g., local adaptation).

To address this idea we develop analytical theory and a set of stochastic simulations. Our analytical theory provides a quantitative description of the problem, but in so doing, we neglect genetic drift (because of the well-known problems with spatial coalescence; Felsenstein (1975b); Barton *et al* (2002)). This should provide a good model for stochastic motion of a single lineage (our main focus), but does not model correlations between lineages or non-Markovian dynamics of the lineages. This approach is common to much theoretical work, that studies the reaction-diffusion equations that govern the deterministic, high-density limit (as in Nagylaki 1975), or the discrete analogues (Hanson 1966). There is also substantial work on how clines are affected by genetic drift (Slatkin & Maruyama 1975; Felsenstein 1975a; Nagylaki 1978; Durrett & Zähle 2007; Barton 2008; Polechová & Barton 2011). Our simulation results suggest that although genetic drift strongly affects local noisiness of the system, it does not substantially influence our conclusions at the moderate population densities we examine, perhaps because we study relatively short time-scales. At much lower population densities, however, local extinction and recolonization may have a stronger effect.

There is also substantial work quantifying the barrier to gene flow caused by clines (Barton 1979a, 1986; Barton & Partridge 2000). We aim to complement this work, and to investigate a possible new approach for identifying genomic regions experiencing selection in hybrid zones.

## Methods

### The Model

We consider two isolated populations — labeled species A and species B — that came into contact *T* generations in the past. We will say that species A was initially on the “left” of the zone of contact, which corresponds to spatial positions *x* < 0. After contact, the populations live across continuous geographical space, with random, local dispersal that we take to be Gaussian (although this should not affect the conclusions if the true dispersal distribution is not too fat-tailed).

We model selection through a single, underdominant locus: at this locus, there are two alleles, one corresponding to each of the ancestral populations. Individuals who are heterozygous at this locus produce on average 1 – *s* times as many offspring than either homozygote. Although most selection in hybrid zones is likely more complex than this simple situation, previous work has shown that such models generate clines that are very similar to models of ecological selection or epistatic systems (Kruuk *et al* 1999; Barton & Shpak 2000), because selection on an allele depends on its marginal effect on fitness. (More discussion of this point in the Discussion.)

As in Sedghifar *et al* (2015), we consider where the ancestors of modern-day individuáis fall across geography as we look further back towards the time of first contact. We say that a locus in a sampled individual is of *ancestry A* if it has been inherited from an ancestor of species *A*, i.e., if its lineage traces back to a spatial position *x* < 0 at the time of secondary contact. A block of genome is of ancestry *A* if every locus in it does the same; this occurs if there is no recombination in this block, or if all lineages generated by recombination events in this block trace back to the *A* side. This process, in which recombination events cause such lineages to *branch*, is illustrated in Figure 1. We say that a block of genome is on the *A background* if it is physically linked to an allele of ancestry *A* at the selected locus; if a block of ancestry *A* includes the selected site then it is necessarily on the *A* background. Because the identity of the selected allele determines how selection acts on the haplotype, and because linked alleles can only move between the *A* and the *B* background in heterozygotes, a key factor in these models is the density and fecundity of heterozygotes.

**Figure 1:**
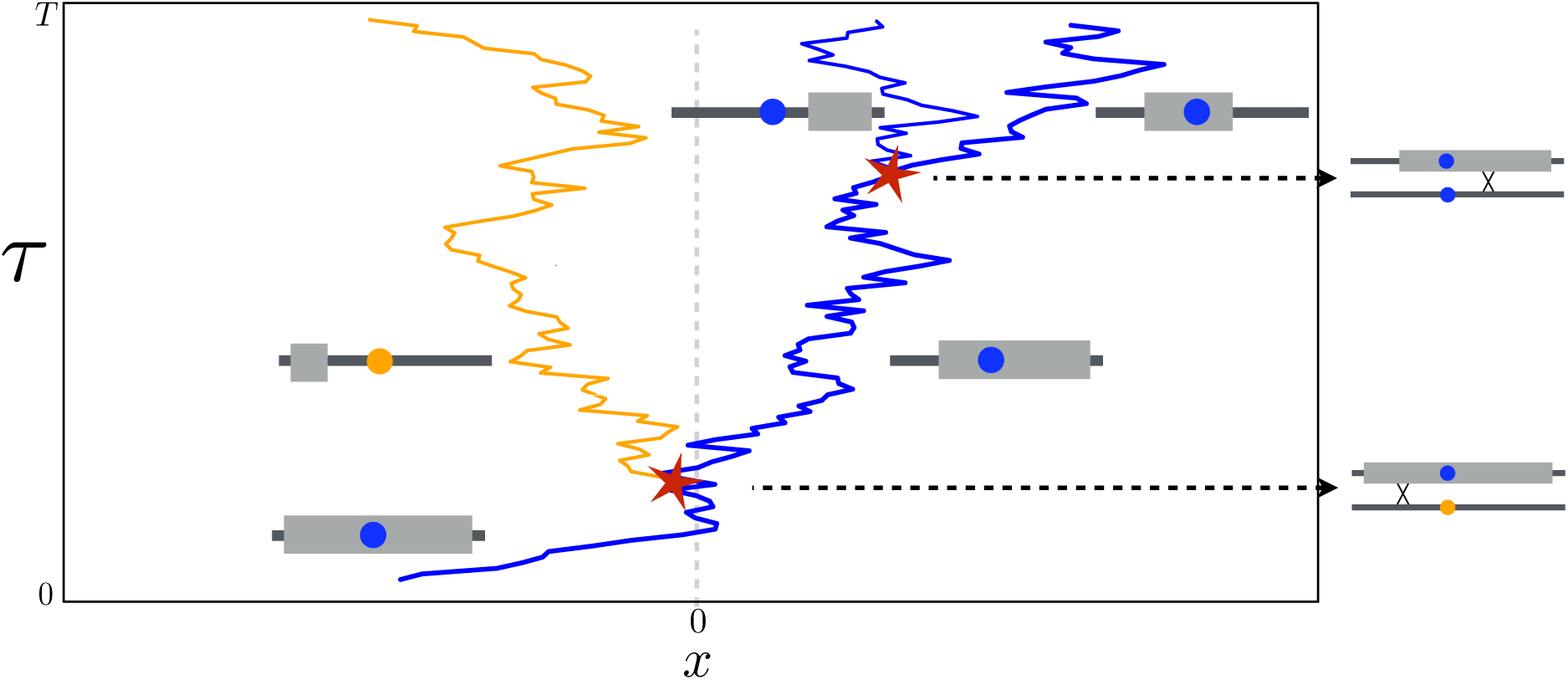
**A schematic figure of the lineages along which a segment of genome has been inherited,** showing the location (horizontal axis) and time (vertical axis) at which the ancestors of a hypothetical sampled haplotype lived. At. the time of sampling (*τ* = 0), the segment contains the *B* allele at the selected locus. Paths traced by lineages across space are depicted by blue and yellow lines. Branching events occur when there was a recombination event. within the sampled segment. The rightmost blue line represents the path of the selected locus. The colors of the other paths indicate the identity at the linked selected locus (blue: linked to allele *B*; yellow: linked to allele *A*). The center of the hybrid zone is at geographic position *x* = 0; in the region *x* < 0, ancestry *B* is less common, and because of selection, the selected lineage spends little time there. Based on position of the lineages at *τ* = 100 generations ago, from left to right the segments are of ancestry (*A*,*B*,*B*).

### Analysis

In this section we aim to give both heuristics for how lineages move, that are helpful in providing intuition and for establishing order-of-magnitude estimates, and analytics, mostly in terms of partial differential equations needing numerical solution.

#### Establishment of the cline at the selected locus

Deterministic theory predicts that a.fter secondary contact, the two alleles at the selected locus will form a stable cline, affecting neighboring loci as well (Barton 1979a; Barton & Bengtsson 1986). First, we turn our attention to how the cline at the selected locus itself is formed. Write *p*(*x*, *t*) for the frequency of the A allele at spatial location *x* and time *t*. Suppose that secondary contact occurred at time *t* = 0, and the A allele is initially fixed on the left, so *p*(*x*, 0) = 1 if *x* < 0 and *p*(*x*, 0) = 0 if *x* > 0. We assume that dispersal is local and unbiased; and that the mean squared displacement between parent and offspring in the direction perpendicular to the cline is *σ*^2^. Assuming that: (i) alleles locally assort into diploids randomly, (ii) habitat and dispersal are homogeneous, and (iii) population density is large, then as in Bazykin (1969), the commonly-used equation that *p* approximately solves is

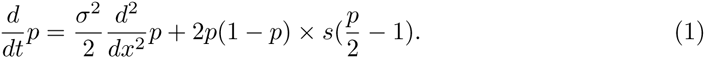

Here the frequency *p* = *p*(*x*, *t*) varies across time and space, and so 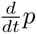 is the local rate of change of the frequency of *A*. The first term, 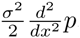, describes the net rate at which *A* alleles arrive by local dispersal. The second term describes the impact of selection on local allele frequency as the product of local genotypic variance, 2*p*(1 – *p*) (equivalently, the local frequency of heterozygotes), and selection on allele *A*, *s*(*p*/2 – 1). The term *s*(*p*/2 – 1) describes underdominance – *A* is favored when common (*p* > 1/2) and disfavored when rare (*p* < 1/2). The equation can be derived as in Nagylaki (1975); the assumption of locally random mating was shown to have only a small effect Christiansen *et al* (1995).

The equation is approximate because it omits terms of order *s*^2^ (so deviations from the prediction occur over 1/*s*^2^ generations), and relies on the Central Limit Theorem to approximate generation-to-generation dispersal by the Gaussian (so if dispersal is non-Gaussian, will fit best over longer time scales, say, tens of generations). As we show below, the solutions provide a good approximation across the range of realistic values of *s*. Although there is no known exact analytic solution to this equation, the steady-state solution is 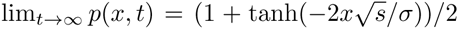 (Bazykin 1969). This relies on several approximations, but the general conclusions should be quite robust: the stable cline has width of order 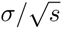, and decays exponentially.

As noted by others (e.g., Slatkin 1973; May *et al* 1975), rescaling space and time by 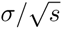 and 1/s respectively in equation (1) results in a dimensionless equation, implying that the cline establishes over a timescale of the order 1/*s*. Since this is how long it takes diffusion at rate *σ*^2^ to smooth across a region of width 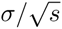, this means we can think of the cline as established by diffusion, despite being slowed down somewhat by selection against heterozygotes.

For simplicity we mostly work with one-dimensional clines, but the mathematics applies as well to two-dimensional systems with some modifications: for instance, if the hybrid zone is a straight line, the description above applies to motion of a lineage transverse to the hybrid zone. In this case, *d*^2^/*dx*^2^ should be replaced by the Laplacian operator. In fact, if the landscape is heterogeneous or dispersal is non-Gaussian, the theory still holds replacing the Laplacian by the appropriate operator. Note, however, that with low population densities, drift can have a strong effect that differs between one and two dimensional systems (Cox & Durrett 1995; Barton *et al* 2010; Durrett & Zähle 2007).

#### Lineages at the selected locus

Now suppose that the frequency profile of the selected allele, *p*(*x*, *t*), is known as a function of time and space. We seek to first understand how lineages of sampled individuals behave at the selected locus, and next extend the analysis to lineages at loci linked to the selected locus.

Consider the collection of *A* alleles found at geographic location *x*. The expected number, among these alleles, that had a parent at location *y* in the previous generation is proportional to the number of *A* alleles that were at *y* multiplied by the per-generation probability of dispersing from *y* to *x*. In other words, lineages move as a random walk determined by the dispersal kernel, but biased towards locations where the selected allele they carry is at higher frequency.

Making the same assumptions as for equation (1), the derivation in Appendix A shows that the lineage of an *A* allele at position *x* moves as Brownian motion driven by dispersal with mean displacement proportional to the spatial derivative of log *p*. This has the effect of pulling *A* lineages towards regions where they are more common, i.e., the left side of the range. (The analogous physical system is a particle that moves at speed *σ*^2^ in the potential – log *p*). To see why this is true, note that the probability that the parent of an *A* allele found at *x* lived at position *x* – *r* is proportional to ℙ{*R* = *r*} *p*(*x* – *r*), where *R* is the random dispersal distance; and so the mean displacement from offspring to parent is **E**[*Rp*(*x* – *R*)]/**E**[*p*(*x* – *R*)]. Using the fact that **E**[*R*] = 0 and **E**[*R*^2^] = *σ*^2^ and expanding *p*() to first order about *x* shows that this mean displacement is approximately *σ*^2^*p*′(*x*)/*p*(*x*), which is *σ*^2^ multiplied by the gradient of log *p*(*x*). This description holds even when the frequency profile of the *A* allele changes with time (replacing *p*(*x*) by *p*(*x*, *t*)).

At equilibrium, the A allele is nearly fixed at a geographic position far to the left (*p* ≈ 1), and is rare far to the right, with mean frequency at distance *x* proportional to 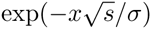. Therefore, roughly, lineages on “their own” side wander randomly, while lineages on “the wrong” side are pushed at constant speed 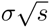 back towards the side where they are more common (since here 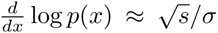 and the speed is *σ*^2^). Since an A allele must, by definition, have been inherited from the A side of the barrier at the time of secondary contact, this push must get more intense the closer it is to the time of secondary contact.

This description gives more information than the steady-state cline, which depends only on 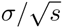. Here we see that lineages with *σ* = 10 and *s* = .16 move much faster than lineages with *σ* = 1 and *s* = .0016, reflecting strong differences in selection against heterozygotes, even though the stable clines have the same form. Even though a rescaling of space and time as described above can make the models equivalent, this difference in lineage speed can be seen through the action of recombination, which we explore next.

#### Lineages at linked loci

The behavior of a lineage at a linked locus is similar to a selected locus. However, there is one important difference – in heterozygotes the lineage linked to the selected locus may recombine onto the other selected background. Therefore, if we follow back through time a lineage at a locus linked to an *A* allele, it will first tend to be inherited from ancestors to the left (as *A* lineages drift to the left). However, with sufficient time in the hybrid zone, recombination allows this linked locus to have been inherited from a *B*-carrying individual, whose ancestors will tend to be more from the right.

Suppose we sample an allele today *r* Morgans from the selected site, and follow its lineage back through time, using *τ* to denote “generations ago” (reserving *t* for time measured in the forward direction). If *X*_*τ*_ is the geographic location of its ancestor *τ* generations ago, then we say that *X* moves as a diffusion pushed by either log(*p*) or log(*1* – *p*) (as described for a selected allele in the previous section). Following this lineage back in time, the identity of the selected allele that ancestor carried at time *τ*, *Z*_*τ*_, jumps between *A* and *B* with each recombination event that occurs between the focal locus and the selected locus (with frequency *r*), that results in a change in the selected background. Thus, by the assumption of locally random assortment of alleles, *Z* shifts from *A* to *B* at rate *r*(1 – *p*), and *B* to *A* at rate *rp* (see Appendix A for a more precise description).

We can describe this Markov process formally in Itô notation: with *B*_*τ*_ a standard Brownian motion, *T*_*B*_(*τ*) the most recent time before *τ* that *Z*_*τ*_ = *B*, and likewise for *T*_*A*_(*τ*),

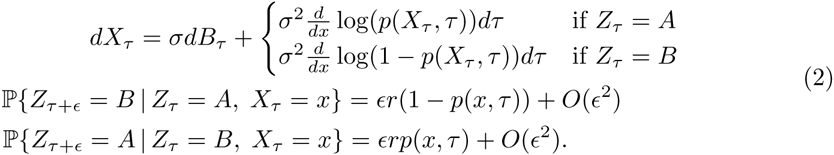

In the first expression, giving the distribution of how the lineage location changes, the first term (*σdB*_*τ*_) is Brownian noise driven by dispersal, and the second is the mean displacement, which moves the lineage “downhill” towards its selected allele’s ancestral range on either – log*p* or – log(1 – *p*), depending on which selected allele the lineage is linked to. (In two dimensions, *d*/*dx* is replaced by the gradient.) The second expression says that the probability a lineage on the *A* background at *x* recombines onto the *B* background is equal to the recombination rate, *r*, multiplied by the local proportion of *B* alleles, (1 – *p*(*x*, *τ*)), per generation.

#### Linked clines

We can use this diffusion model for lineages to find expected clines in ancestry, i.e., the expected proportion of individuals who inherit from species *A*, as a function of space, time, and position on the genome. Precisely, we need the probability that an allele sampled *t* generations after secondary contact at location *x*, at recombination distance *r* to a selected allele of type *z*, is inherited from an individual of ancestry *A*, where *z* can be either *A* or *B*. We denote this probability *q*_*z*_(*x*, *t*, *r*). In the notation above,

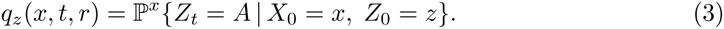

(Recall that *Z* depends implicitly on *r*.) The description above implies that *q*_*z*_ solves the following Kolmogorov backward equation:

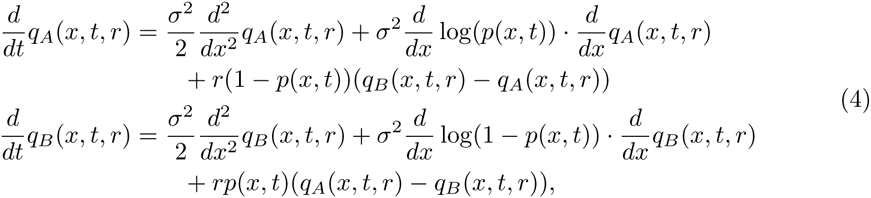

with boundary conditions *q*_*A*_(*x*, 0, *r*) = 1 and *q*_*B*_(*x*, 0, *r*) = 0. The three terms in these equations come from Brownian movement of a linages (i.e., the smoothing action of local dispersal), the tendency of a lineage to inherit from regions where its type is more common (i.e., the net flux induced by reduced fitness in the hybrid zone), and recombination between selected backgrounds, respectively. We note that because recombination occurs within a deme, the recombination terms in equation (4) are quite standard (Hartl & Clark 1989).

We will have more use for the differential operators on the right-hand sides of these equations, so define these as 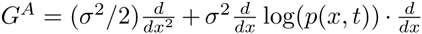 and 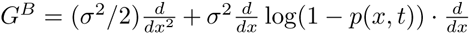, so that equation (4) can be written more compactly as

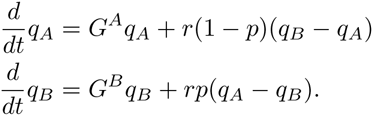

In technical terms, *G*^*A*^ and *G*^*B*^ are the generators of the diffusions of lineages of selected alleles of ancestry *A* and *B*, respectively; informally, they encode the stochastic motion of lineages at the selected loci.

#### Numerical computation

To determine how the cline at a linked locus is expected to relax, we can solve the partial differential equations (1) and (4) numerically. For instance, Figure 2 shows a heatmap of *p*(*x*, *t*, *r*), the expected frequency of ancestry *A* at location *x* and time t at a site at recombination distance r from the selected site, which is computed as the frequency of ancestry *A* on each selected background weighted by the frequencies of each background: *q*(*x*, *t*, *r*) = *p*(*x*, *t*)*q*_*A*_(*x*, *t*, *r*) + (1 – *p*(*x*, *t*))*q*_*B*_(*x*, *t*, *r*). The equations are solved numerically in R, using the ReacTran package (Soetaert & Meysman 2012).

**Figure 2:**
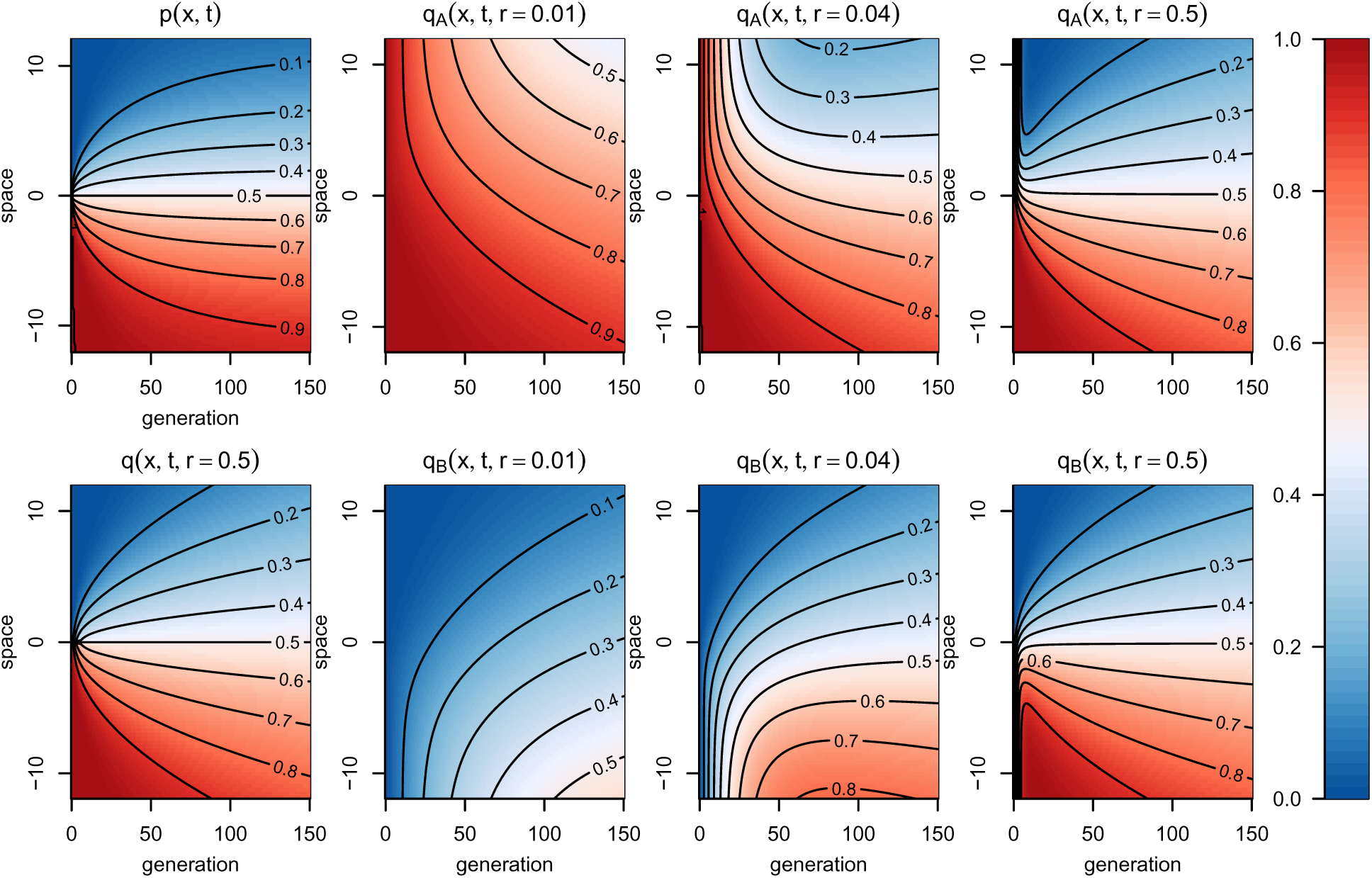
**Probabilities of A ancestry,** across space (vertical axis, in units of *σ*) and time (horizontal axis, in generations). In each plot, color corresponds to the expected frequency of *A* ancestry at a particular location in time and space. The selection coefficient is *s* = .02. **Top left:** at the selected site, showing establishment and stabilization of the cline on a time scale of 1/*s* = 50 generations. **Bottom left:** at an unliked site, with cline flattening continuing with 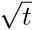. **Remaining figures** show frequencies of *A* ancestry *conditional* on the ancestry at the selected site, at different distances from the selected site (*r* = .01, .04, and 0.5 Morgans), as described in the text (see definition of *q*_*z*_(*x*, *t*, *r*)). See Figure S1 for the same figure over a longer period of time.

#### Haplotype lengths

We now develop our model further to find the frequency at which *entire* blocks of genome (haplotypes) of a single ancestry are found, at a given location and time. Suppose we sample an individual at spatial location *x* and time *t* after the initiation of gene flow, and genotype them on the genomic segment between positions *a* and *b*, relative to the selected site. We are interested in the probability *g*_*z*_(*x*, *t*; *a*, *b*) of finding an entire segment (*a*, *b*) of ancestry *A* given that the individual has selected allele of type *z*. For instance, *g*_*B*_(0, 10; 0, .2) is the probability that an individual carrying a selected *B* allele sampled at the center of the zone at *t* = 10 has a block of *A* ancestry for 0.2 Morgans to the right of the selected site.

As in Sedghifar *et al* (2015), a given block of genome is inherited along a single lineage ever since the most recent recombination event that fell within that block. Prior to this, there are two lineages to follow (see Figure 1), and so lineages behave as labeled, branching diffusions, where the total branching rate is conserved. We do not consider subsequent coalescence. The general description of the process, again looking backwards in time, is as follows: The lineage of a segment of genome moves as a linked locus described in equations (2), with recombination distance r equal to the rate at which the segment recombines away from the selected site. (If the selected site lies inside the segment, *r* = 0.) Additionally, at rate equal to the genetic length of the segment, recombination occurs within the segment, at which point it splits in two at a uniformly chosen location between *a* and *b*, each of which proceeds as before, independently. In this description, *g*_*z*_(*x*, *T*; *a*, *b*) is then the probability that all branches are found on the *A* side of the hybrid zone at the time of secondary contact *T* units of time ago. We will write *r*(*a*, *b*) for the distance from the segment to the selected site: always taking *a* ≤ *b*, *r*(*a*, *b*) = 0 if *a* ≤ 0 ≤ b and *r*(*a*, *b*) = min(|*a*|, |*b*|) otherwise.

The resulting equation is similar to (4) with a term added for branching, which is written (omitting the (*x*, *t*) for conciseness):

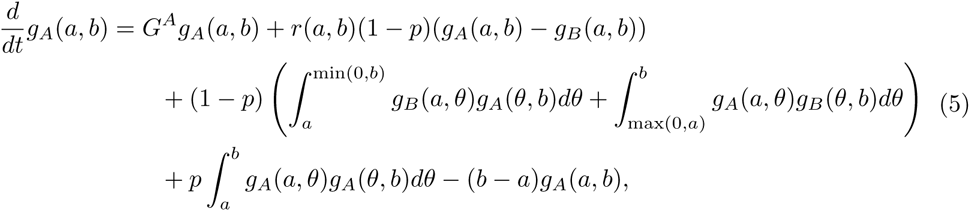

Here the first term (*G*^*A*^*g*_*A*_) represents spatial mixing, and the second term results from recombinations *between* the block and the selected site in heterozygotes, which switch the identity of the linked, selected locus without splitting the block (the factor (1 – *p*) is the probability that the block, initially linked to an *A* allele, encounters a *B* allele under the assumption of locally random mating). In the integrals, *θ* is the genomic position of the recombination event. The third term accounts for such recombinations in a heterozygote: the portion of the block nearest the selected site remains linked to an *A* allele, and the remaining portion becomes linked to a *B* allele (e.g., the lower recombination event shown in Fig. 1). To account for cases where the selected locus lies outside the interval [a,b], we say that the integral 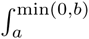 is zero if *a* > 0 and likewise 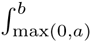 is zero if *b* < 0. The last integral results from recombination inside the block in homozygotes for *A* (e.g., the upper recombination event in Fig. 1), and the final term balances the outflux due to all recombinations. The equation for *g*_*B*_(*x*, *t*; *a*, *b*) is identical after exchanging *A* ↔ *B* and *p* ↔ (1 – *p*). The boundary conditions are *g*_*A*_(*x*, 0; *a*, *b*) = 1 and *g*_*B*_ (*x*, 0; *a*, *b*) = 0.

#### Numerical solutions

Notice that equations (5) are hierarchical in (*a*, *b*): the equation for haplotype identity probabilities on a segment (*a*, *b*) depends only on those probabilities for segments contained in (*a*, *b*). This is useful for numerical solutions, described in more detail in Appendix B.

#### Correlations in ancestry

To compute correlations in local ancestry (i.e., “ancestry disequilibrium”, as in Pool (2015); Schumer & Brandvain (2016)), we need only follow lineages at two sites, instead of an entire region. Doing so only requires computing correlations in ancestry between markers, which can be done directly using our numerical code; see Appendix B for more detail.

#### Simulations

We implemented forwards-time simulations of a one-dimensional grid of demes with non-overlapping generations and fixed population sizes (a Wright-Fisher model). Individuals are diploid, with haploid number *n* = 2 chromosomes, each of length 1 Morgan. One chromosome pair harbors, at position 0.5M, a single locus that reduces fitness by *s* in heterozygotes, while the other contains no sites under direct selection.

Each deme has exactly *N*_*d*_ diploid hermaphroditic individuals at the start of each generation. Then, every individual disperses to a (possibly) new deme by choosing a Gaussian displacement with mean zero and variance *σ*^2^, then dispersing to the nearest deme. (The mean displacement is zero, and when comparing to theory we compute *σ* as the standard deviation of this discrete distribution.) Migrants past either end of the range remain at the terminal demes. Then, fitnesses are computed (heterozygotes at the selected locus have fitness 1 – *s*; homozygotes have fitness 1), and in each deme *N*_*d*_ pairs of parents are chosen, with replacement, with probability proportional to their fitness and selfing allowed. Since migration is not conservative, demes may have no available parents; in this case, parents are chosen from other demes with probability proportional to fitness multiplied by exp(–*x*^2^/*σ*^2^), where *x* is the distance to the other deme. The next generation is formed by carrying out meiosis in each parent and combining the resulting gametes such that each pair of parents leaves one descendant. Meiosis results in alternating blocks of the gamete’s chromosome being inherited from the two parental chromosomes, with the blocks separated by a Poisson(1) number of uniformly chosen recombination points along the chromosome, and the order of the parental chromosomes chosen randomly. The simulation software works by recording, for each chromosome, a list of ancestry breakpoints, and the index of the ancestor from which the chromosome inherited that genomic region. We then assigned ancestry at individual loci by looking up which side of the zone the ancestor lived on.

For efficiency, in most simulations the number of demes was only 3–5 times wider than the stable cline width. We verified that the size of the simulated region did not affect dynamics in the center of the zone by running additional simulations on wider regions (as we did for numerical solutions of PDE); an example is shown in Figure S5.

The simulations were executed in R, with scripts available at https://github.com/petrelharp/clinal-lineages.

#### Measures of introgression

The above theory and simulations generate predictions about patterns of ancestry surrounding selected loci. In reality, however, such loci are not usually known, so it is useful to have per-site statistics that may allow for detection of candidate targets of selection. The most straightforward measure is *l*_*B*_(*m*, *x*), the mean length of all contiguous segments of ancestry *B* sampled at position *x* that contain genomic position *m*. Likewise, *l*_*A*_(*m*, *x*) is the mean length of segments of ancestry *A*. We compute similarly the unconditioned mean block length *l*(*m*, *x*) by averaging all segment lengths without regard to ancestral identity.

We also look at the mean length of the two chunks, m – and m+, that flank, to the left and right respectively, the block of unbroken ancestry containing m. As described below, these blocks tend to be *shorter* than average when m is the selected site, motivating us to define the statistic

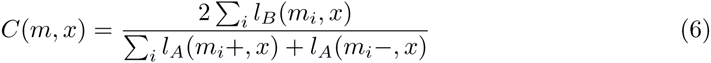

where *m*_*i*_ is the block length in individual *i*, and the sum is over individuals at location *x*. The statistic *C* is the mean block length around m in the population, divided by the mean lengths of the two blocks directly flanking the block containing *m*.

To identify regions of the genome with abnormal distributions of block lengths, we compute *l*_*B*_(*m*, *x*) and *C*(*m*, *x*) at a grid of positions across geography and across the genome. From these, we compute normalized versions *l*̅_*B*_(*m*, *x*) and *C*̅(*m*, *x*) by dividing by the empirical mean across the genome for each geographic location *x*: for instance, *l*̅_*B*_(*m*, *x*) = *n*_*x*_*l*_*B*_(*m*, *x*)/Σ_*i*_ *l*_*B*_ (*m*_*i*_, *x*), where the sum is over the *n*_*x*_ individuals at location x. While there is useful information contained in the geographic distribution of block length patterns, obtaining good spatial sampling can often be difficult, and this allows us to search for patterns if block lengths are only known within relatively few populations.

We are able to partially trace the genealogy of haplotypes in our simulations. In particular, we wish to know the relative number of ancestors from *T* generations ago that are represented in present day populations at a given locus, given the frequency of a particular ancestry. This is a reflection of the average size of a haplotype’s family, and is potentially a source of additional information about selection. Within each deme *x*, we calculated *F*_*B*_(*m*, *x*), which is the number of independent ancestors from time *T* contributing to the pool of alleles of ancestry *B* at site *m*, divided by the number of individuals of ancestry *B* at site *m*.

## Results

### Single locus clines

As expected, clines in simulations form and begin to flatten, with ancestry frequencies at the selected site maintaining a stable shape, and tightly linked sites flattening more slowly than distant sites. This is seen in Figure 3, and supplementary Figures S2, S3 and S4. Clines in ancestry frequency matched expected values found from deterministic theory up to stochasticity due to genetic drift, which is more pronounced at lower population densities. For instance, Figure 3 shows that expected clines computed from equation (4) match simulated clines quite well. Figures S16, S17, and S18 show this comparison with a lower population density than in Figure S2, and separating linked clines based on the identity of the linked selected allele. The agreement is good over hundreds of generations, although as the clines flatten they are seen to wobble (which happens at rate proportional to 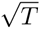; Barton (1979b)).

**Figure 3:**
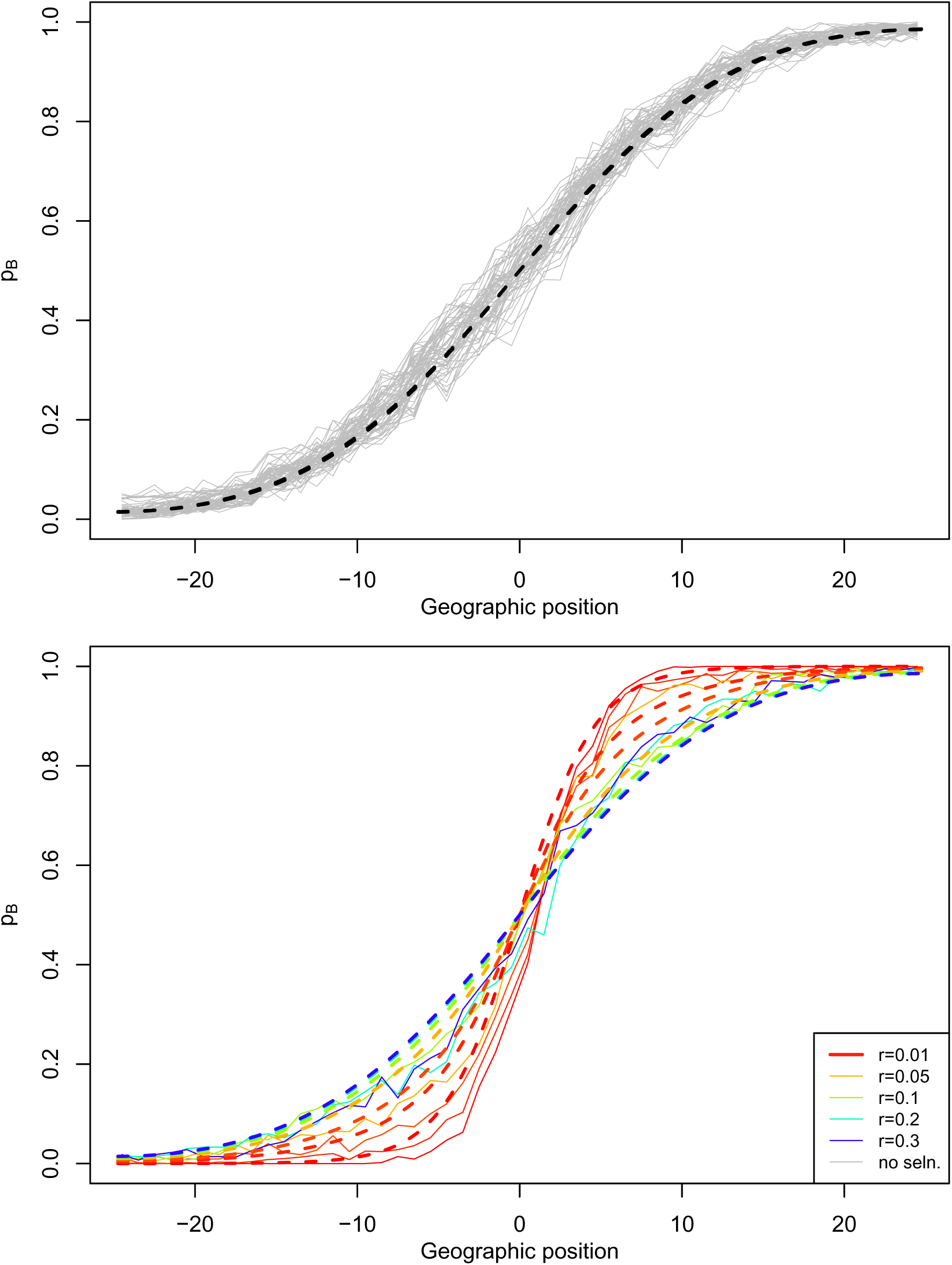
**Comparison of simulated to theoretical clines:** Frequency of ancestry *B*, across geography at different physical positions on the genome, simulated for a hybrid zone *T* = 100 generations after secondary contact, with *s* = 0.1, using 50 demes, each with 500 diploid individuals and *σ* = 1. In each figure, each line represents a locus some distance *r* away from the true target of selection, and *r* = 0 represents the locus that is under selection. Dotted lines are expected frequencies of *B* ancestry 1 – *q*(*x*, *t*, *r*) (unconditional on the ancestry of the linked allele), computed numerically. **Top:** loci across a chromosome unlinked to the selected locus. **Bottom:** loci across the chromosome carrying the selected locus, with colors corresponding to different values of r, from red (tight linkage to selected site) to blue (distantly linked). The two sets of clines are shown superimposed in Figure S2.

Loci not under direct selection can in principle spread across the cline unimpeded. In practice, however, it can take quite some time for even unlinked neutral loci to homogenize, due to the decreased fitness of heterozygotes (Barton & Bengtsson 1986) and the relative slowness of diffusive movement. We display the spread of ancestry across space and time in Figure 2. The cline in ancestry at a locus r Morgans from the selected site will have flattened out to distance *x* (say, on the *B* side) if there is a good chance that the corresponding lineage that begins linked to a B allele traces back to an *A* allele on the opposite side of the hybrid zone. Since lineages linked to *B* alleles move nearly as unbiased Brownian motion on the *B* side, this is only possible if Brownian motion has had enough time to travel distance *x*, i.e., if 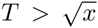. This square-root flattening is seen in Figure 2 (and is discussed for environmentally determined clines by May *et al* (1975); but see Durrett & Zähle (2007)). A linked lineage must also spend at least 1/*r* generations in heterozygotes to have a good chance of recombining, so clines with *r* < 1/*T* will still resemble the selected cline, which can also be seen in Figure 2.

In principle, the genomic window about the selected site in which clines remain narrow could be quite a bit wider, since the only way to move linked lineages between selected backgrounds is via recombination in a heterozygote, and heterozygotes for the selected allele are only found at high frequency in the cline. The majority of lineages are generally pushed away from the cline but have no bias far away, so the amount of time a lineage spends in heterozygotes should grow as 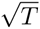 for large *T*, and so the width of the genomic region showing clines about the selected locus could be substantially larger than 1/*T*. However, this distinction appears hard to observe for realistic parameter values.

### Blocks of ancestry

The distribution of contiguous ancestry block lengths contains more information than allele frequency alone. We are specifically interested in how the tracts of ancestry surrounding the selected locus compares to the rest of the genome. Ideal information – true ancestry assignments for a few simulated individuals sampled from across space – are shown in Figure S6 (*T* = 100 generations) and Figure 4 (*T* = 1000 generations). For the more recent hybrid zone (*T* = 100), the selected cline has established, but linked clines are still flattening. After a longer period (*T* = 1000), clines over much of the chromosome are flat (since the width of the entire population is less than 1/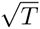 = 31.6*σ*), but a distinct enrichment of each ancestry is observed around the selected site.

**Figure 4:**
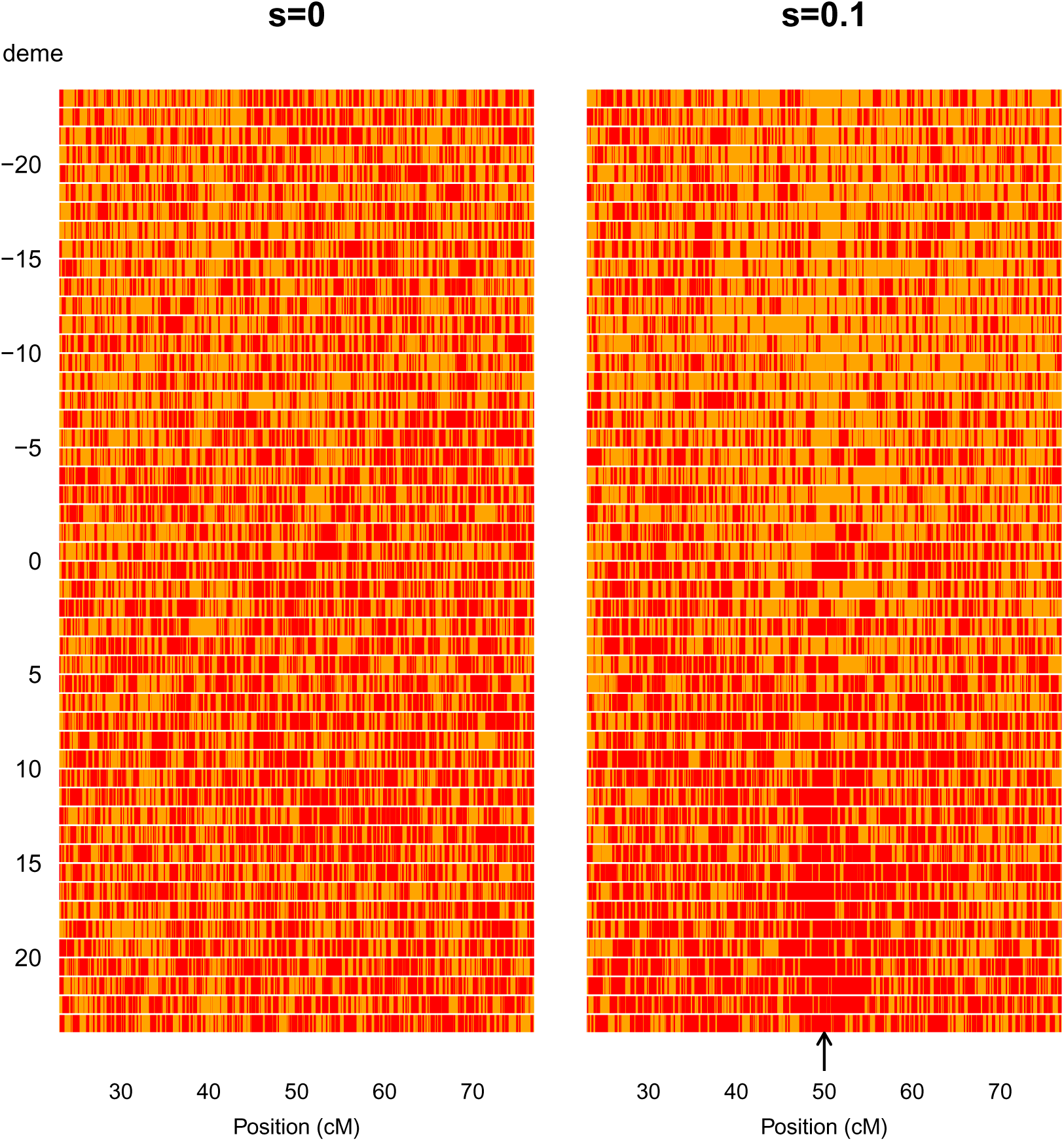
**Ancestry blocks for randomly sampled chromosomes** across a hybrid zone of age *T* = 1000. Here we compare chromosomes of length 1M from a neutral zone to a zone that has a single under-dominant locus with *s* = 0.1 in the middle of the chromosome (indicated by black arrow). Red blocks along the chromosome denote ancestry *B*, and orange blocks are ancestry *A*. The simulation was performed in a population with 50 demes, each with 500 diploids, and *σ* = 1. An analogous figure at *T* = 100 is shown in Figure S6.

We expect that, in the absence of selection, blocks of *A* ancestry across the genome will tend to be shorter the further one goes into the *B* side of the cline, because they have had more opportunities to recombine with *B* haplotypes. This is seen in Figure S8. However, we expect that stretches of *A* ancestry *containing a selected site* will be longer than those that do not contain the selected site at the same spatial location, because lineages containing the selected site have usually been inherited from the *A* side of the cline recently. As discussed above, we expect these lineages move at speed roughly 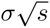, so (selected) A alleles at distance x from the cline center have last had an ancestor on the A side of the cline around 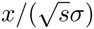 generations ago (compared to *x*^2^/*σ*^2^ for a neutral allele). This implies the scale on which A haplotypes are found surrounding the locus should be no longer than about 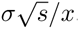.

#### Identifying selected loci

The statistics *l*̅(*m*, *x*) and *C*(*m*, *x*) show promise for identifying selected loci under some circumstances. As expected, regions surrounding a locus under selection are more resistant to introgression, as seen in Figures 4 and S6. When present, we expect haplotypes that contain the locally less common allele to be longer than the genome-wide average. Indeed, as shown in Figures 5, S9 and S10, the mean length of such haplotype blocks is up to three times longer than the average for that geographic location, peaking quite sharply around the location of the selected site.

**Figure 5:**
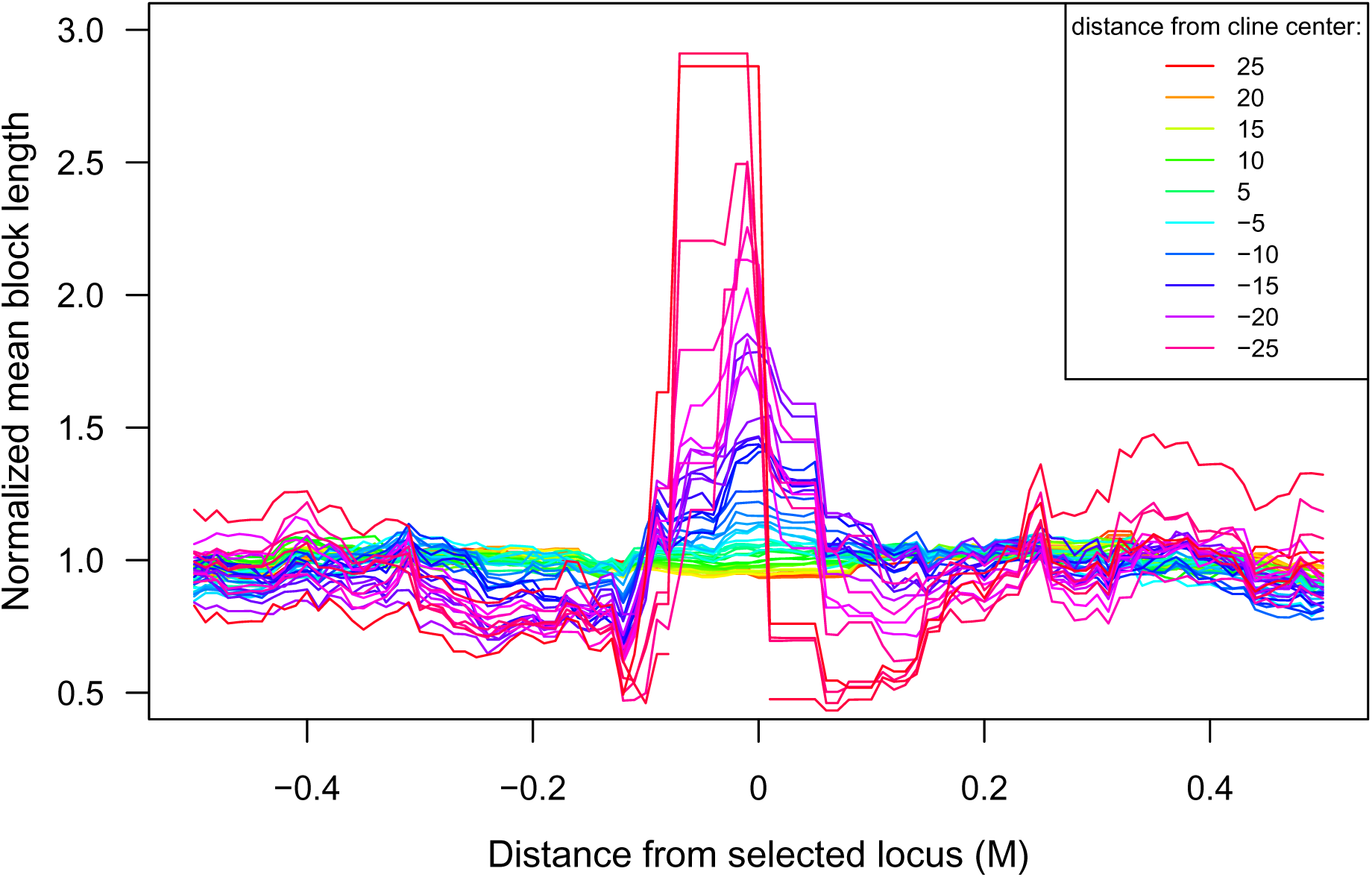
**Normalized mean enclosing block length,** *l*̅_*B*_(*m*, *x*), along a simulated chromosome of length 1*M* with selected locus of *s* = 0.01 at position 0.5*M* in a hybrid zone of *T* = 1000. Here each line shows *l*̅_*B*_(*m*, *x*), in a given deme some distance from the zone center. Chromosome were sampled across the hybrid zone, which consists of 50 demes, each containing 500 diploid individuals. The same simulation and statistic are shown in Fig. S9, on a coarser genomic scale.

Using the fact that genomic regions are inherited from ancestors *T* generations ago in blocks of size roughly 1/*T*, we can provide some intuition about the physical scale of the expected signal. There will be a long segment of *A* ancestry if many such adjacent blocks are all inherited from the *A* side. Ancestries of neighboring blocks are correlated, due to the branching process described above. But, if we assume they are independent, then since the block surrounding location *r* in an individual at geographic location *x* is of ancestry *A* with probability q*A*_*A*_(*x*, *T*, *r*), we’d expect to see, roughly, *q*_*A*_(*x*, *T*, *r*)(1 + 2*q*_*A*_(*x*, *T*, *r*))/(1 – *q*_*A*_(*x*, *T*, *r*)) consecutive blocks of ancestry *A* about a given site unlinked to the selected site. (This assumes, crudely, that the number of A-blocks on either side of the enclosing block has a Geometric distribution with parameter *q*_*A*_(*x*, *T*, *r*).) This implies the mean length *l*(*x*, *r*) would be *q*_*A*_(*x*, *T*, *r*)(1 + 2*q*_*A*_(*x*, *T*, *r*))/((1 – *q*_*A*_(*x*, *T*, *r*))*T*), and the mean conditional length *l*_*A*_(*x*, *r*) would be (1 + 2*q*_*A*_(*x*, *T*, *r*))/(*T*(1 – *q*_*A*_(*x*, *T*, *r*)). Furthermore, we know that if *x* = 0 or if *T* is large and *r* is not too small, that *q*_*A*_ is close to 1/2 (so *l*_*A*_(*x*, *r*) ≈ 4/*T*); and for *r* < 1/*T* that *q*_*A*_ looks like the selected cline. Also, we know from the discussion above that the lineage of a *selected A* allele, if it is in the region where *A* is rare, moves at speed roughly 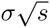 back towards the *A* side of the zone, returning to the region where *A* is common in about 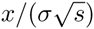 generations. Therefore, a selected *A* allele found on the *B* side of the zone should carry with it a haplotype of average length 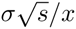 that looks like haplotypes from the center of the zone. This analysis suggests that *A* haplotypes in the center of the zone should be of average length 4/*T*; this is indeed what is seen at distant sites, for instance, in Figure S7. Haplotypes at the selected site are expected to be longer, but still of a length proportional to 1/*T*, suggesting that the normalization in the statistic *C*(*x*, *m*) is appropriate, as shown in Figure S13, although a numerical prediction of the value of *C*(*x*, *m*) is elusive.

The mean haplotype length found without conditioning on ancestry, *l*(*x*, *m*), shows a smaller increase near the selected locus, because most blocks will be of the locally common type, and so do not trace back to regions of different block lengths (Figures S8, S12 and S11).

Power will be optimal at intermediate values of *s*. If selection is too strong, it may be difficult to observe this signal due to a lack of introgressed selected sites, while if selection is weak, the selected lineage does not move very fast, and the hybrid zone is wider, allowing more recombination, and so the strength of signal from elevated *l*_*B*_(*m*) is diminished. For instance, no signal is seen in Figure S19. For similar reasons, power and resolution are best at intermediate *T*.

### The size of migrant families

Within-ancestry haplotype diversity, i.e., the number of ancestral haplotypes of each type, could provide additional information about whether introgression is through relatively few, successful migrants, or through many migrants that each contribute relatively little. In our simulations, the average local family size of a selected *B* allele (*F*_*B*_(*m*, *x*)) decreases with distance into the *A* side of the hybrid zone, and is lowest far away from the zone center, where ancestry *B* is at low frequency (Figure 6). Unlinked loci have local family sizes similar to neutral simulations, and loci linked to the selected locus have intermediate sizes. This pattern is consistent with the prediction that unfit lineages tend to be recent migrants, which will have smaller families.

**Figure 6:**
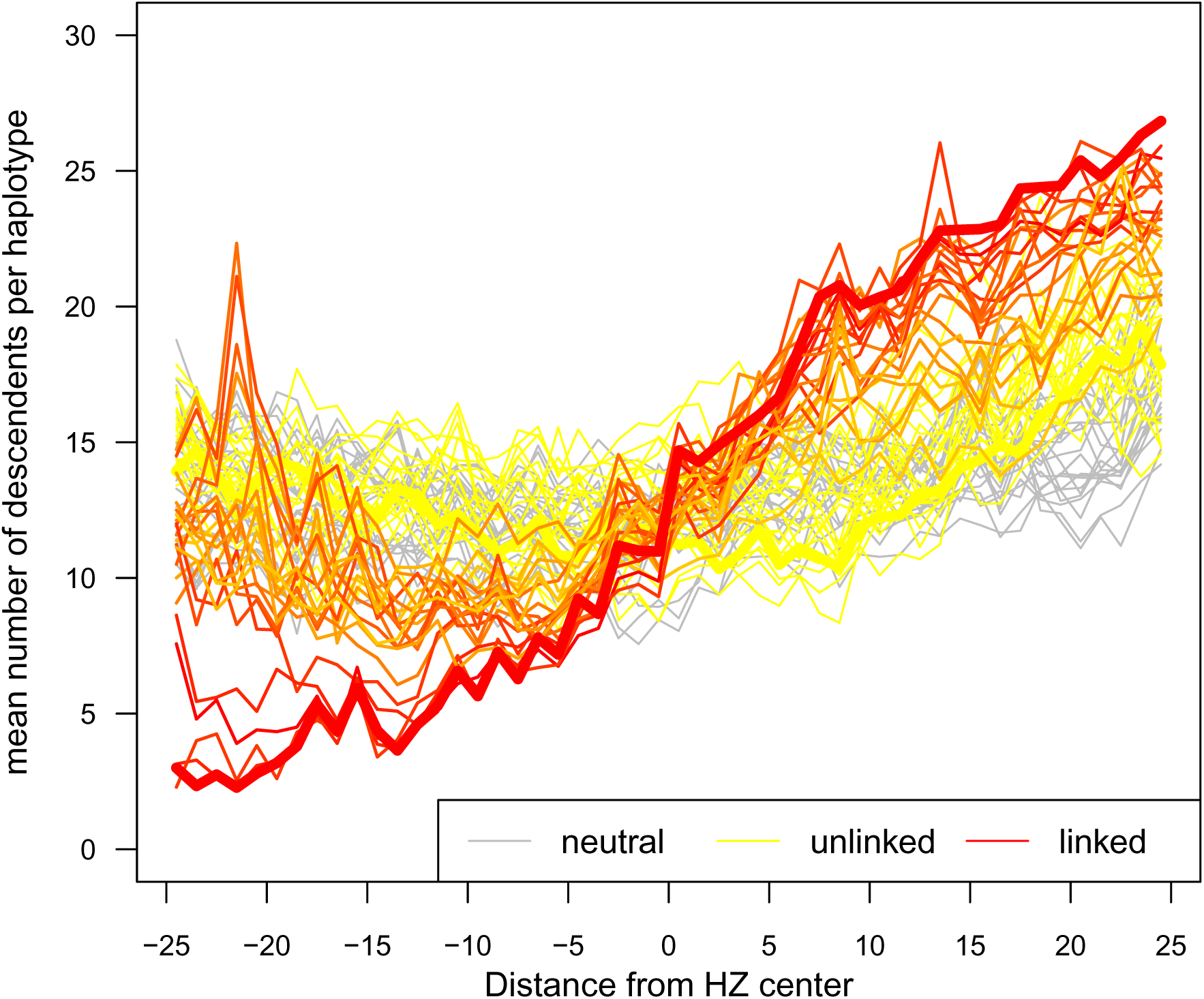
**Mean family size of haplotypes.** The number of individuals of ancestry *B* per number of ancestors (*F*_*B*_(*m*, *x*)) from secondary contact occurring *T* = 1000 generations ago, represented across geographic space. The hybrid zone is 50 demes, each containing 500 diploid individuals. Each red or orange line represents a site some distance away (ranging from 0 – 0.5cM) from the selected site (here *s* = 0.01 and *σ* = 1). Yellow lines are corresponding positions on an unlinked chromosome with no selected loci, and grey lines are corresponding positions in a simulation with no selection. Bold lines depict the target of selection when present, and corresponding position on chromosomes not harboring any selected sites.

## Discussion

Using a combination of theory and simulations, we present a description of the process of cline formation and haplotype structure in a relatively young (i.e., non-equilibrium) hybrid zone. We show why clines establish over time 1/*s*, and why lineages of selected loci tend to move back towards their ‘ancestral home’ when in a geographic region where they are unfit. This occurs at speed 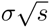. Based on this we predict, and observe in simulations, that blocks of ancestry surrounding these selected loci are longer, especially when distant from the center of the cline, than those surrounding neutral loci. This extends previous theoretical work on hybrid zones, which has primarily focused on stable clines in allele frequency. Additionally, our work suggests that the ancestry block length distribution can help detect targets of purifying selection in hybrid zones. The resolution of this approach is expected to scale with 1/*T*, as this is the physical scale over which linked clines persist.

### Genomic signals associated with targets of selection

Popular approaches to identifying loci under selection in hybrid zones involve identifying alleles that are exceptional in terms of frequency across space, or genome-wide admixture proportion (Porter *et al* 1997; Gompert *et al* 2012). The availability of genomic data has made it possible to use local ancestry as an additional source of information. In particular, ancestry deconvolution facilitated by programs such as hapMIX (Price *et al* 2009), LAMP (Sankararaman *et al* 2008) and fineSTRUCTURE (Lawson *et al* 2012) can inform the demographic history of hybridization/admixture from present day samples (e.g., Hellenthal *et al* 2014). We described how selection against hybrid incompatibilities results in long contiguous blocks of ancestry around these loci. This is because regions surrounding selected loci do not readily introgress, and so introgressed alleles have been inherited from “their” side of the zone recently. We get quantitative understanding from the fact that lineages of loci under selection move as Brownian motion pushed at speed 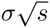 towards its ancestral side. In other words, disadvantageous alleles that have encroached deep into the other side of the hybrid zone have done so by chance and, since they do not persist for long on the “wrong” side, are likely to have done so relatively recently. Haplotypes surrounding the selected locus therefore have had relatively few opportunities for recombination with different ancestries and this is reflected in longer blocks of contiguous ancestry.

Through our simulations we find also that blocks of ancestry that have crossed the hybrid zone and are closely linked to the selected site without overlapping it (i.e., blocks that are adjacent to the block containing the selected site) are shorter than average for the spatial location (Figure S13). An intuitive explanation for this pattern is that loci physically linked to a selected site have recently come from the center of the zone or beyond. Compared to other chromosomes in their new geographic location, these migrants will have on average longer *B* haplotypes at the selected locus, and shorter *A* haplotypes nearby.

Our results suggest that the statistic *l*_*B*_(*x*, *m*) – the mean genetic length of ancestry surrounding a site of a given ancestry – could help identify selected loci in hybrid zones. The ratio of mean adjacent block lengths, *C*(*x*, *m*), also shows a very sharp peak at the selected site, suggesting that, despite the fact that haplotypes surrounding incompatibilities might be quite long, genome scans may have the power to extract fine-grained regions near selected loci from the large chunks of ancestry that will often flank these regions.

Our results focus on the length distribution of ancestry blocks — a statistic that can only be obtained in the few systems for which dense genetic markers and phase information is known. However, as the cost of sequencing continues to drop and long-phased reads become more common, the patterns described here could further aid in the identification of selected sites in hybrid zones in many non-model taxa. In the meantime, patterns of elevated pairwise LD, which is easily computed from readily available low-density and unphased genomic data, could offer an alternative path forward for empirical work. Importantly, the numerical solutions derived above readily and efficiently predict correlations in ancestry, and so our results can be applied immediately to genotype data for any taxa with markers placed on a genetic map.

Ancestry assignments have the additional benefit that relatively unbiased estimates are possible even with markers of problematic ascertainment such as those on SNP arrays. The power and resolution of these approaches depends strongly on the strength of selection, the time since secondary contact, and the strength of genetic drift: in our simulations, we found good power and resolution at *s* = .01, *T* = 1000 generations after secondary contact, and with hundreds of individuals per dispersal distance.

### Assumptions

Here we review the assumptions we made and their likely impact.

#### The nature of selection

Although our assumed scenario of selection against heterozygotes at a single locus is uncommon in nature, theoretical and simulation studies show that clines formed by underdominant selection and extrinsic selection are effectively indistinguishable without additional information (May *et al* 19T5; Barton & Gale 1993; Kruuk *et al* 1999), especially over short time scales (Barton 1979b), and their shape is determined mostly by the mean fitness of the heterozygote (Slatkin 1973).

Our assumption of a single selected locus may have greater consequences. Our model is most relevant to scenarios with few targets of selection scattered throughout the genome, and therefore, our predictions may differ significantly from situations in which the density of selected sites is higher. Having numerous selected sites within one ancestry block increases the strength of selection (Barton & Bengtsson 1986), and the relatively short map distances between linked incompatibilities generates a longer unit which is not readily broken up by recombination (Barton 1986). Both factors are expected to result in regions that are surrounded by even longer segments of unbroken ancestry, and will impact expectations of genome-wide pattens of clines and block lengths, as well as the resolution to which one could detect targets of selection (Slatkin 1975; Barton 1983).

#### Geography

To speed calculations and simulations, we assumed a homogeneous, one-dimensional geographical range. Our analytical results further assumed a large population density, effectively working with a deterministic model that ignores coalescence and associated stochasticity. In contrast, our simulations model regularly spaced demes of finite size. In reality, populations may be patchily connected, especially at the edges of species ranges where hybrid zones may occur. The degree to which inhomogeneous geography would affect the predictions depends on how patchy the zone is; the differential equations provide a way to evaluate this in specific circumstances.

#### Coalescence

Extending our analytical results to capture stochasticity arising from coalescence/pedigree structure represents an important future direction. Indeed, correlated fluctuations visible in simulations (e.g., Figure S9) are likely due to coupling due to demographic stochasticity; and simulations at lower density show larger fluctuations than those at higher density. Furthermore, ignoring pedigree structure can result in an underestimate of covariance in ancestry, as we have ignored additional sharing of ancestry through shared genealogy (Liang & Nielsen 2014); but there is nonetheless good agreement between analytic predictions and simulations (which include an explicit pedigree structure). It could be particularly important for applications to understand to what degree drift within an old, stable hybrid zone can mimic the haplotype patterns seen in a recently formed zone, in the same way that drift can produce clines at neutral loci in old hybrid zones.

#### Theory and simulation

We have taken two complementary approaches, using both simulation and theory, and comparing the two. As usual, simulations make fewer biological simplifications, while theory provides more generalizable conclusions. To do this, we have described the branching diffusion process that approximates the lineages along which haplotypes are inherited. Since the expected motion of a lineage depends on the local frequencies of the selected alleles, these diffusions are time-inhomogeneous.

The diffusion model for lineages predicts that quantities of interest solve sets of coupled partial differential equations (PDE), which we have written down. As there are no known analytical solutions to these PDE, we have constructed numerical solutions (and provide the source code for doing this). A main role of these solutions in our work has been to verify that theory based on the diffusion model of lineages matches realistic, individual-based models. These solutions easily and quickly provide predictions of joint frequencies at small numbers of loci: about 1 second to compute predicted clines, as opposed to hours for the full simulation. However, due to the high dimensionality of the haplotype problem (spatial position × time × endpoints of the haplotype), numerical solutions for mean haplotype lengths along the genome can be as computationally intensive as simulations (although are substantially less noisy). More work could be done to develop more efficient methods of solution, but it may be better to perform more biologically realistic forwards-time simulations that includes coalescence and drift to better characterize the block length distribution. If however the correlation in ancestry, rather than the block-length distribution, is of interest the PDE approach may be preferable because it is easily modified to provide predictions for spatial and temporally inhomogeneous systems – for instance, across maps of real landscapes.

#### Patterns of divergence

A number of studies have described heterogeneous patterns of genetic divergence across the genome. Work on these “islands of divergence” (Turner *et al* 2005; Nosil *et al* 2009) and related patterns have been largely descriptive (Cruickshank & Hahn 2014; Noor & Bennett 2009). Our study here contributes to a model-based understanding of how migration and selection may influence such patterns across the genome of hybridizing populations. Overall, focusing on lengths of ancestry blocks across the genome brings focus to the processes of migration and selection rather than high-level summaries that are somewhat abstracted from the evolutionary process.

#### Adaptive introgression

While our focus has been on hybrid incompatibilities, unconditionally adaptive loci are expected to easily introgress across hybrid zones (Barton 1979a; Barton & Bengtsson 1986; Martinsen *et al* 2001; Arnold 2004). Future work could take a similar approach to understand how positive selection shapes ancestry block lengths, and predict signatures of adaptive introgression in hybridizing populations using similar statistics presented here. These could eventually be combined to gain a fuller understanding of the forces shaping patterns of introgression in hybrid zones. In particular, beneficial alleles tightly linked to incompatibilities cannot introgress until recombination separates them; our model provides some rough expectation on how quickly this should happen.

## Acknowledgements

We thank Graham Coop, Nancy Chen, Emily Josephs, Kristin Lee, Simon Aeschbacher, Jeremy Berg, Vince Buffalo, Nick Barton and two anonymous reviewers for helpful feedback and suggestions. This work was supported by National Institute of Food and Agriculture (NIFA) project #1007008 to YB and by National Science Foundation grant DBI-1262645 and a Sloan Research Fellowship to PR.

## Data Accessibility

All scripts used in production of this paper are available at https://github.com/petrelharp/clinal-lineages under the GPLv3 software license.

## A Rescaling a discrete model to obtain the lineage motion

For concreteness, here we describe a discrete model that rescales to the continuous model we consider.

In this discrete model, the total number of individuals at location *x* is *N*(*x*), and we count “individuals” as *haploid*, so each individual is of type either *A* or *B* at the selected locus, and the proportion of individuals of type *A* and location *x* and time *t* is *p*(*x*, *t*) (but, we often neglect the *t*). Suppose that type *A* individuals at location *x* reproduce at rate *s*_*A*_(*x*), and likewise type B at rate *s*_*B*_(*x*). Assuming locally random mating, we will then have that *s*_*A*_(*x*) = 1 – *s*(1 – *p*(*x*)) and *s*_*B*_(*x*) = 1 – *sp*(*x*).

At reproduction, individuals recombine with others in the same location, with recombination occuring between the locus we follow and the selected locus with probability *r*, and the offspring choose a new location *y* with probability *m*(*x*, *y*). The population dynamics are random, but suppose that *N*(*x*) is sufficiently large that these do not vary substantially with time. Suppose this is a Moran model. There are four things that can happen:

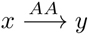 One type A individual at location *x* reproduces, either does not recombine or recombines with another type A, and sends the offspring to *y*.
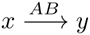 An individual at location *x* reproduces, recombines with the other type, and sends to *y* an offspring who inherits at the selected locus from the type *B* parent and the at the neutral locus from the type A parent.
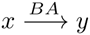 An individual at location *x* reproduces, recombines with the other type, and sends to *y* an offspring who inherits at the selected locus from the type A parent and the at the neutral locus from the type *B* parent.
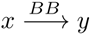 One type *B* individual at location *x* reproduces, either does not recombine or recombines with another type *B*, and sends the offspring to *y*.

These four things happen at rates:

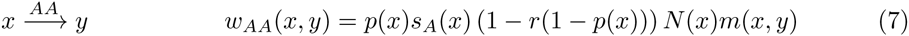

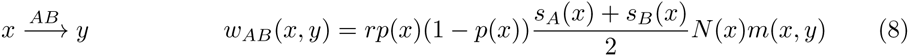

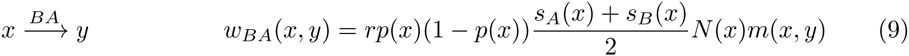

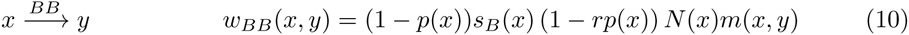

For instance, there are *p*(*x*)*N*(*x*) type *A* individuals at *x*, that reproduce at rate *s*_*A*_(*x*); the chance that each reproduction is with a type *B* is (1 – *p*(*x*)), and the chance there is a recombination that gives the offspring genotype *AB* is *r*/2; the offspring has probability *m*(*x*, *y*) to disperse to *y*; the total rate at which individuals at *x* with the *A* allele produce offspring at *y* with the *B* allele is the product of these terms, *p*(*x*)*N*(*x*)*s*_*A*_(*x*)(1 – *p*(*x*))*r*/2. The second and third rates (for 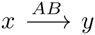 and 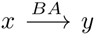) are the same; they are this value plus the corresponding term for individuals carrying the *B* allele producing offspring that carry the *A* allele.

Note that at equilibrium, we require *N* to solve

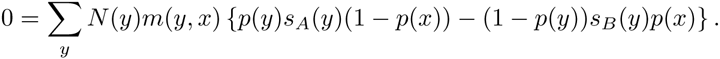

### Lineage movement

These rates tell us the rates at which a lineage will move, backwards in time. For instance, the rate at which a lineage at the selected locus currently in a type A individual at location *x* jumps to another type A individual at location *y* is equal to the rate of influx of migrants from *y* divided by the number of A alleles at *x*, or

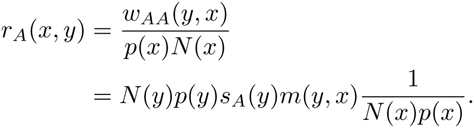

Let *X*_*t*_ denote the position of the lineage of a selected locus of time *A* at time t in the past, and let *f* be test function with *f*(*ρ*) = 0. Then,

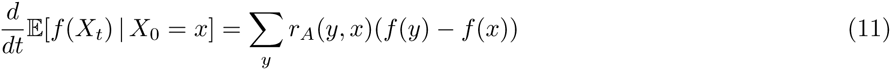

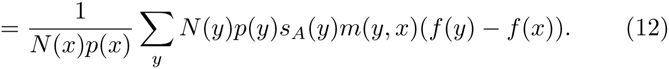

### Diffusion limit

Now suppose that *m*(*x*, *y*) is symmetric, and depends on a parameter *σ* so that as *σ* → 0, the associated random walk converges to Brownian motion, so that for an arbitrary smooth function *f*,

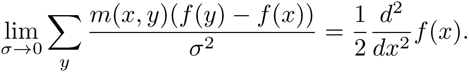

Write 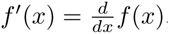, and note that

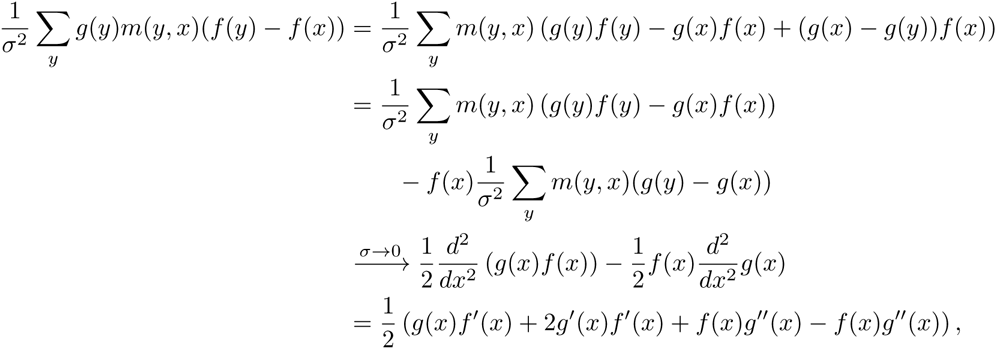

which tells us the differential operator that best approximates the discrete sum:

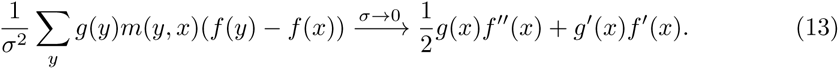

Under these assumptions, combining (12) and (13),

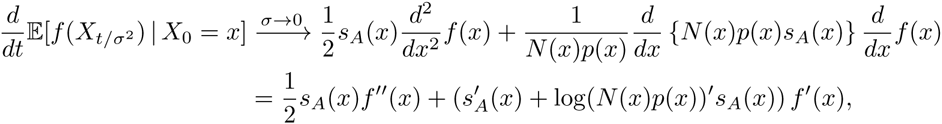

i.e. *X*_*t*/*σ*^2^_ converges to a diffusion with mean displacement (“drift” in diffusion terminology) 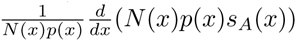 and killed at rate *ρk*(*x*).

In our case, since *s*_*A*_(*x*) = 1 – *s*(1 – *p*(*x*)), the drift is *p*′(*x*)/*p*(*x*) to first order in *s*; the time scaling by *σ*^2^ implies that the Brownian noise and the mean displacement should both be scaled by *σ*^2^.

## B Numerical calculation of haplotype probabilities

In this section, we give details for how we found numerical solutions to the partial differential equations (PDE) of the text, which are all of reaction-diffusion type. The R code, with worked examples, is available in our git repository. Spatial grids were usually chosen to have at least four grid sites per dispersal distance, but using substantially finer grids did not affect the results.

The forwards-time evolution of the selected alleles, equation (1), presents no difficulty; we use the ReacTran package (Soetaert & Meysman 2012) to compute discrete approximations to the diffusion term, and the deSolve package (Soetaert *et al* 2010) to solve the equation.

The equations (4) describing probabilities that a lineage descends from A ancestry, conditional on the linked selected allele, required some more attention. First, *s* > 0.1 the system of equations can be *stiff* (as is commonly observed for reaction-diffusion equations), and hence slow to solve, because of the extremely steep slope of the selected allele frequency *p*(*x*, *t*). In practice we used *s* < 0.1. Second, the ReacTran function tran.1D that converts the diffusion portion of the PDE into a system of ODE contains a term like

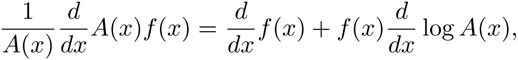

where *A*(*x*) is the interface area between grid cells, and *f*(*x*) is the flux. (See Soetaert & Meysman (2012) for more details.) Discrete approximations of the left-hand side, as implemented in ReacTran, run into numerical difficulties if *A*(*x*) is small, and in our case, *A*(*x*) is equal to *p*, the local frequency of the selected *A* allele. To avoid this, we made minor modifications to the tran.1D to provide a discrete approximation to the right-hand side.

Haplotype probabilities are obtained from equations (5), which is a coupled system of integro-differential equations in three variables plus time. One method for solution would be via a Wild sum over the number of recombination events, as we did in Sedghifar *et al* (2015). Here, we solved the equations numerically, again discretizing the equations and using the ode.1D function of the deSolve package. The reaction-diffusion part is the same as for equations (4). The functions *g*_*A*_(*x*, *t*; *a*, *b*) and (*x*, *t*; *a*, *b*) are functions of space (*x*), time (*t*), and the endpoints of the block in question (*a* and *b*, with *a* < *b*). The second integral in (5) is

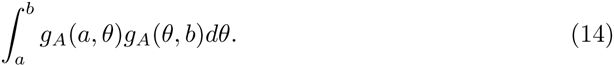

Conceptually, this is an integral transformation 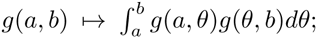 the reason it appears here is that for the entire segment (*a*, *b*) to be of ancestry *A*, if there was a recombination at *θ*, the two segments (*a*, *θ*) and (*θ*, *b*) must both be of ancestry *A* (and, ignoring coalescence, these probabilities are independent). Suppose we have divided the segment of chromosome into a regular grid, say, *r*_1_ < … < *r*_*n*_. The natural discretization approximates this transformation by a sum, and is equivalent to keeping track of only a finite number of loci. Writing *g*_*A*_(*i*,*j*) for *g*_*A*_(*x*, *i*; *r*_*i*_, *r*_*j*_), the discrete transformation we use corresponding to (14) is *g*_*A*_(*i*, *j*) ↦ the sum

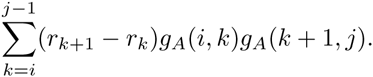

This is the correct term for the process only tracking loci on the grid, because for all the alleles at *r*_*i*_, *r*_*i* + 1_,…, *r*_*j*_ to be of ancestry *A*, when a recombination occurrs between *r*_*k*_ and *r*_*k* + 1_, the sequences of alleles at *r*_*i*_,…, *r*_*k*_ and at *r*_*k* + 1_,…, *r*_*j*_ must each be of ancestry *A*. The first integral term in (5) is

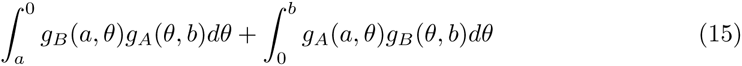

We now require that one of the grid points along the chromosome is exactly at the selected site; say this is *r*_*ℓ*_ = 0. The discrete term corresponding to (15) is

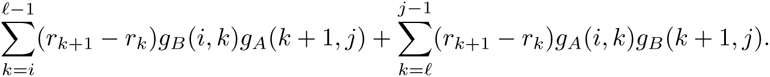

Since these are not easily vectorizable in R, for efficiency we implemented the discrete transformations in C, using the Rcpp package (Eddelbuettel & Francois 2011). In implementing these, we kept track of *g*_*A*_(*i*, *j*) in a vector in the order that the upper triangular elements of a matrix are encountered when traversing the matrix column-wise, allowing for efficient computation of the sum.

### A note on symmetry

The equations we present have the symmetry that they are invariant after exchanging *A* and *B* and reversing space. For instance, *q*_*A*_(*x*, *t*, *r*) = 1 – (–*x*, *t*, *r*). Using this fact would speed up the code by a factor of two, at the cost of generality: as written, it would be easy to modify the code to allow space to be inhomogeneous (which would break this symmetry).

## C Supplementary Figures

**Figure S1:**
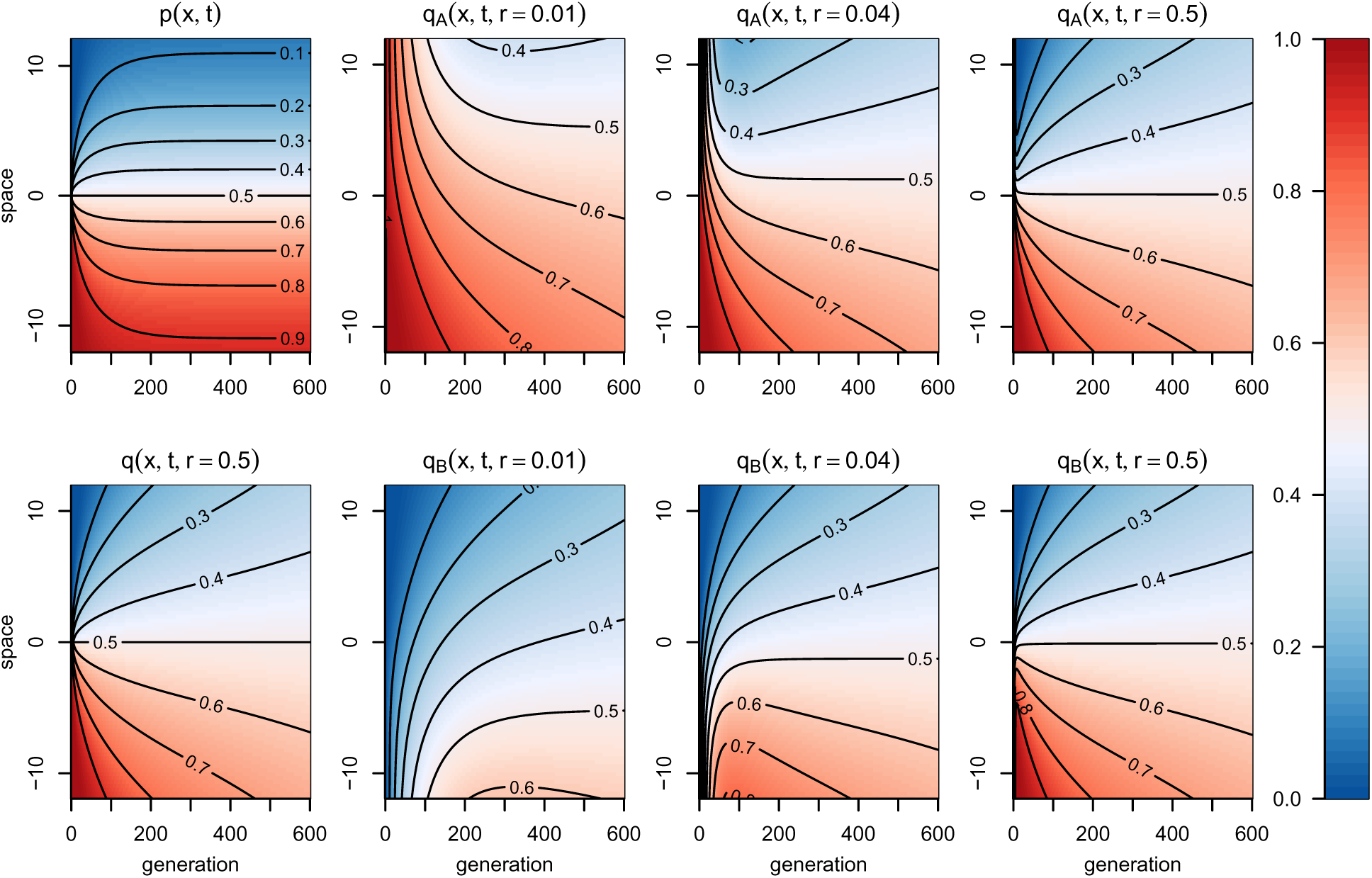
**Probabilities of *A* ancestry,** across space (vertical axis, in units of *σ*) and time (horizontal axis, in generations). In each plot, color corresponds to the expected frequency of *A* ancestry at a particular location in time and space. The selection coefficient is *s* = .02. **Top left:** at the selected site, showing establishment and stabilization of the cline on a time scale of 1/*s* = 50 generations. **Bottom left:** at an unliked site, with cline flattening continuing with 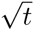. Remaining figures show frequencies of *A* ancestry *conditional* on the ancestry at the selected site, at different distances from the selected site (*r* = .01, .04, and 0.5 Morgans), as described in the text (see definition of *q*_*z*_(*x*, *t*, *r*)). See figure 2 for the same figure over a shorter period of time.

**Figure S2:**
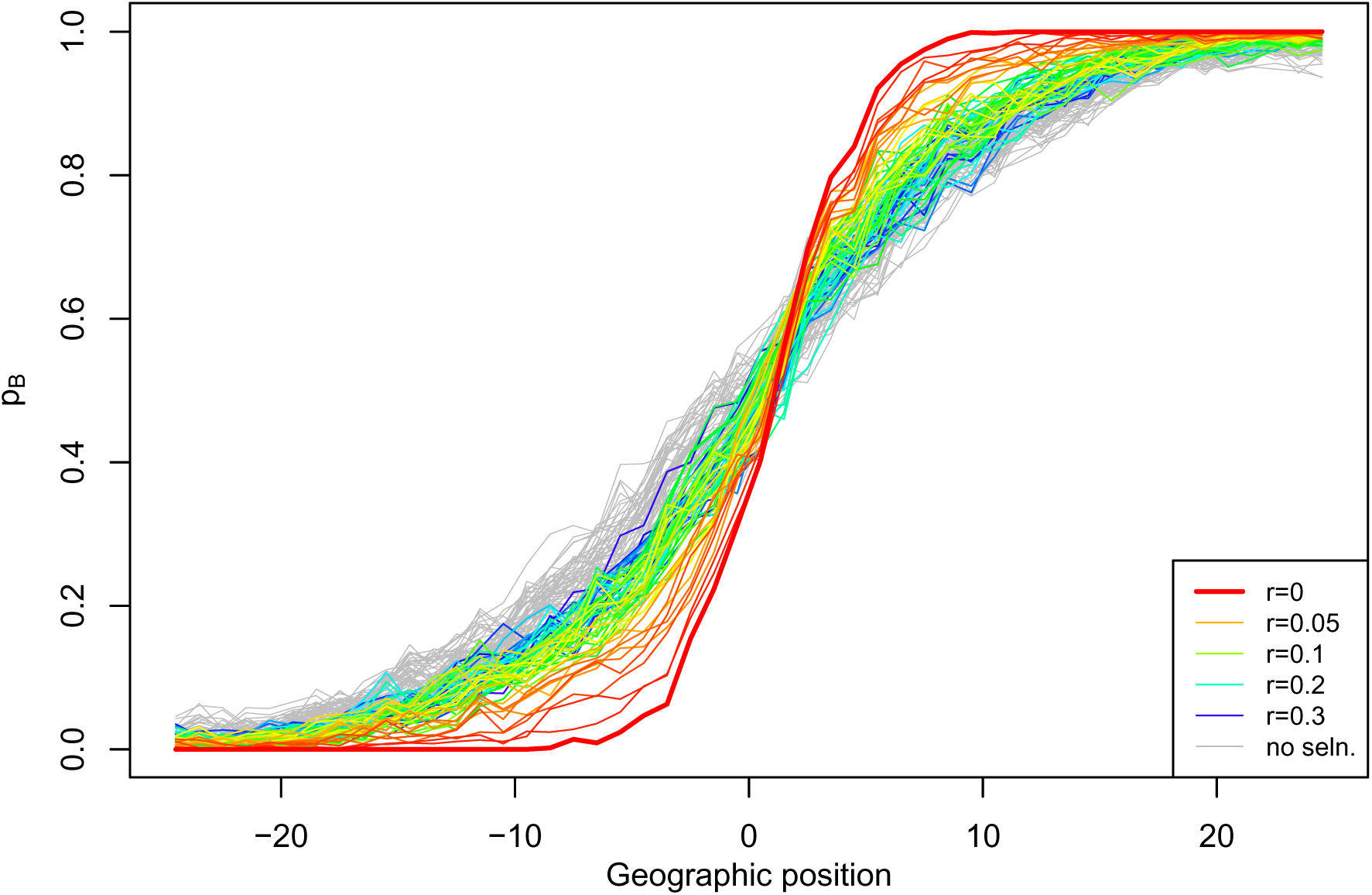
Frequency of ancestry *B*, across geography at different physical positions on the genome, simulated for a hybrid zone *T* = 100 generations after secondary contact, with *s* = 0.1, using 50 demes, each with 500 diploid individuals and *σ* = 1. Each line represents a locus some distance r away from the true target of selection with colors corresponding to different values of r, transitioning from red (tight linkage to selected site) to blue (distantly linked). Grey lines represent the same positions from a simulation with identical parameters except that *s* = 0. Corresponding theoretical quantities are shown juxtaposed in Figure 3; the same plot is shown with weaker selection in Figure S3 and at a longer time in Figure S4.

**Figure S3:**
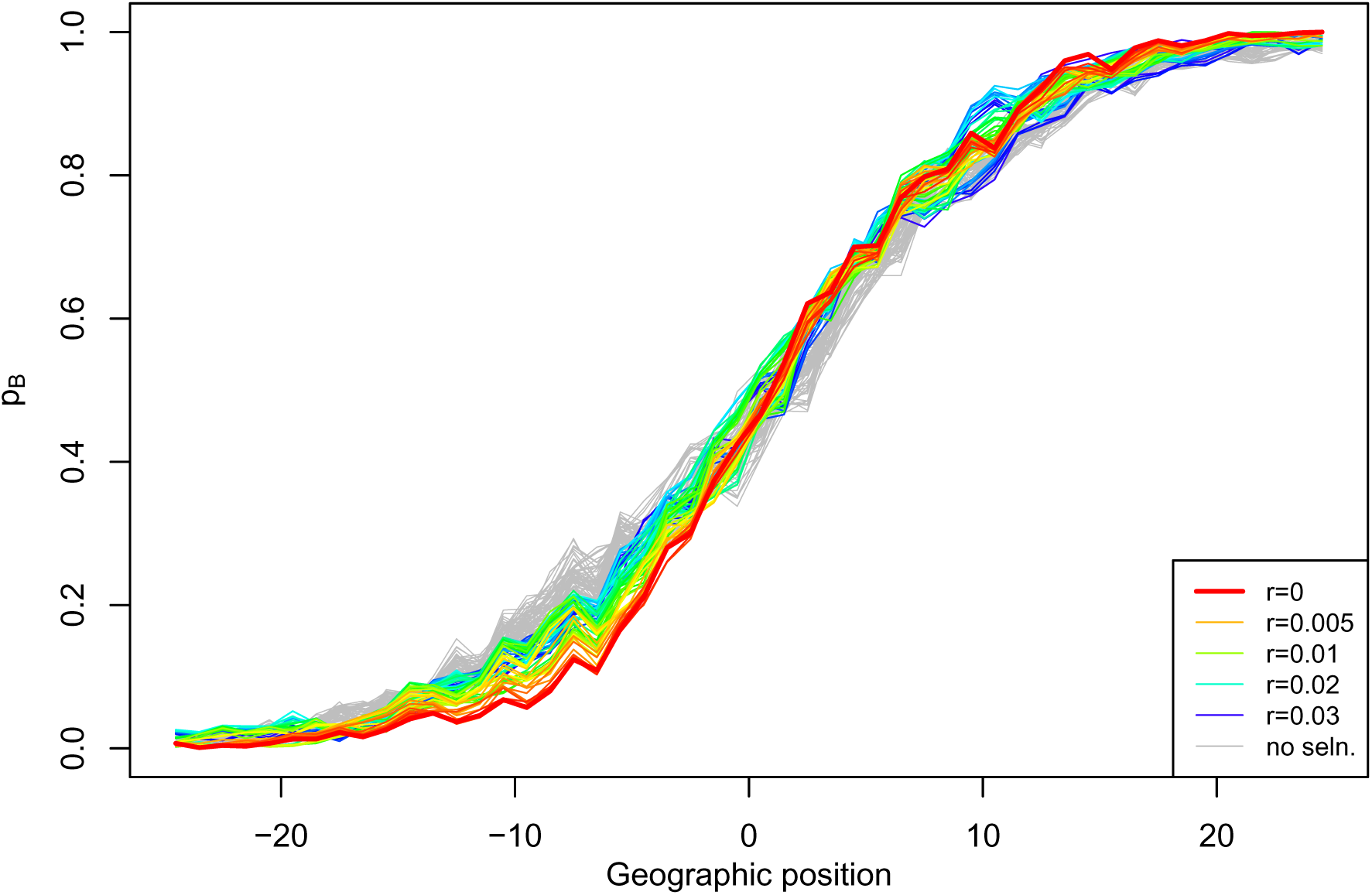
Frequency of ancestry *B*, across geography and at several physical positions on the genome, simulated for a hybrid zone *T* = 100 generations after secondary contact, and with *s* = 0.01. The simulated zone had 50 demes, each with a population size of 500 diploid individuals. Each line represents a locus some distance *r* away from the true target of selection, and *r* = 0 represents the locus that is under selection. Grey lines represent the same positions from a simulation with identical parameters except that *s* = 0.

**Figure S4:**
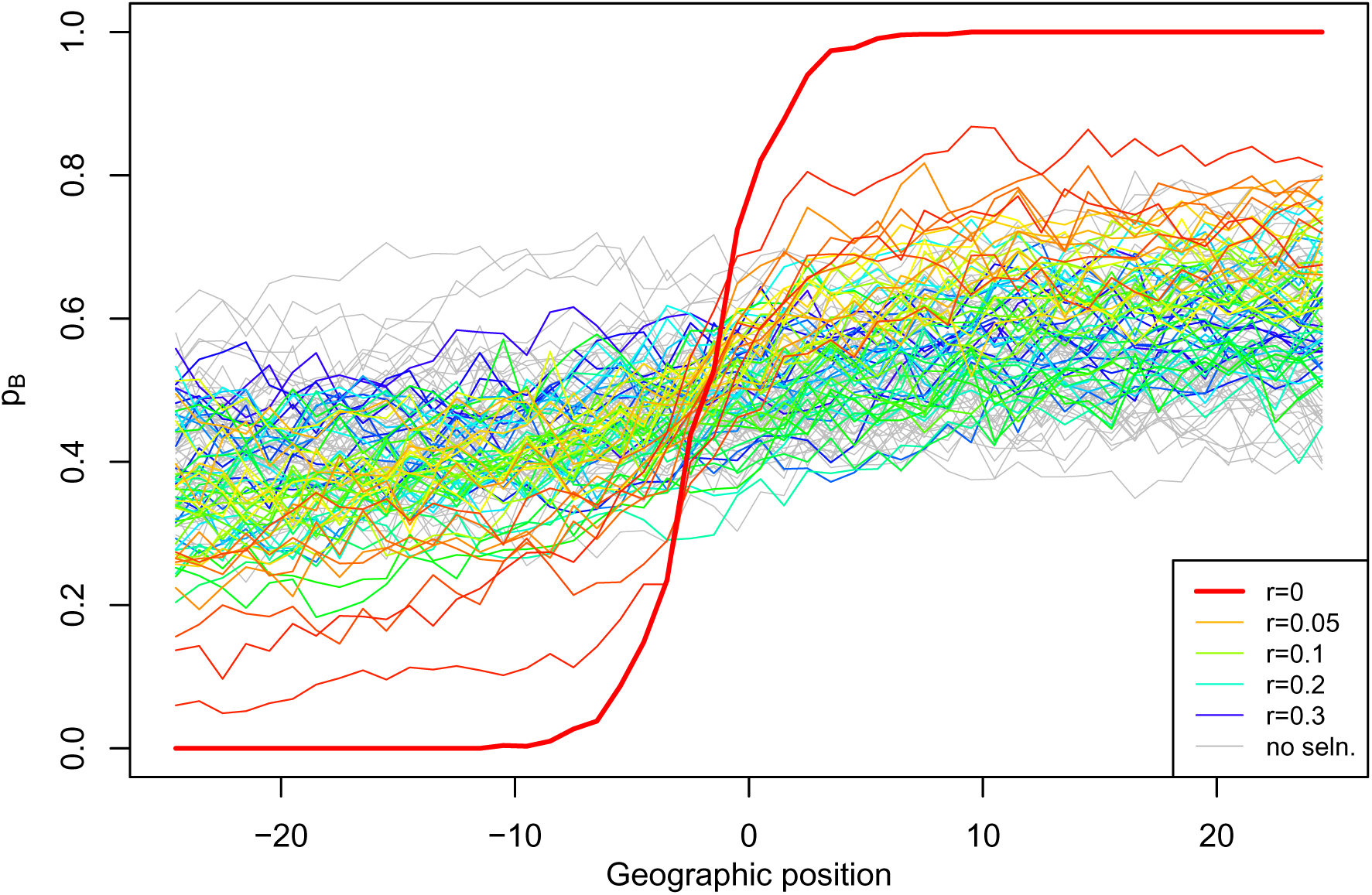
Frequency of ancestry *B*, across geography and at several physical positions on the genome, simulated for a hybrid zone *T* = 1000 generations after secondary contact, and with *s* = 0.1. The simulated zone had 50 demes, each with a population size of 500 diploid individuals. Each line represents a locus some distance r away from the true target of selection, and *r* = 0 represents the locus that is under selection. Grey lines represent the same positions from a simulation with identical parameters except that *s* = 0.

**Figure S5:**
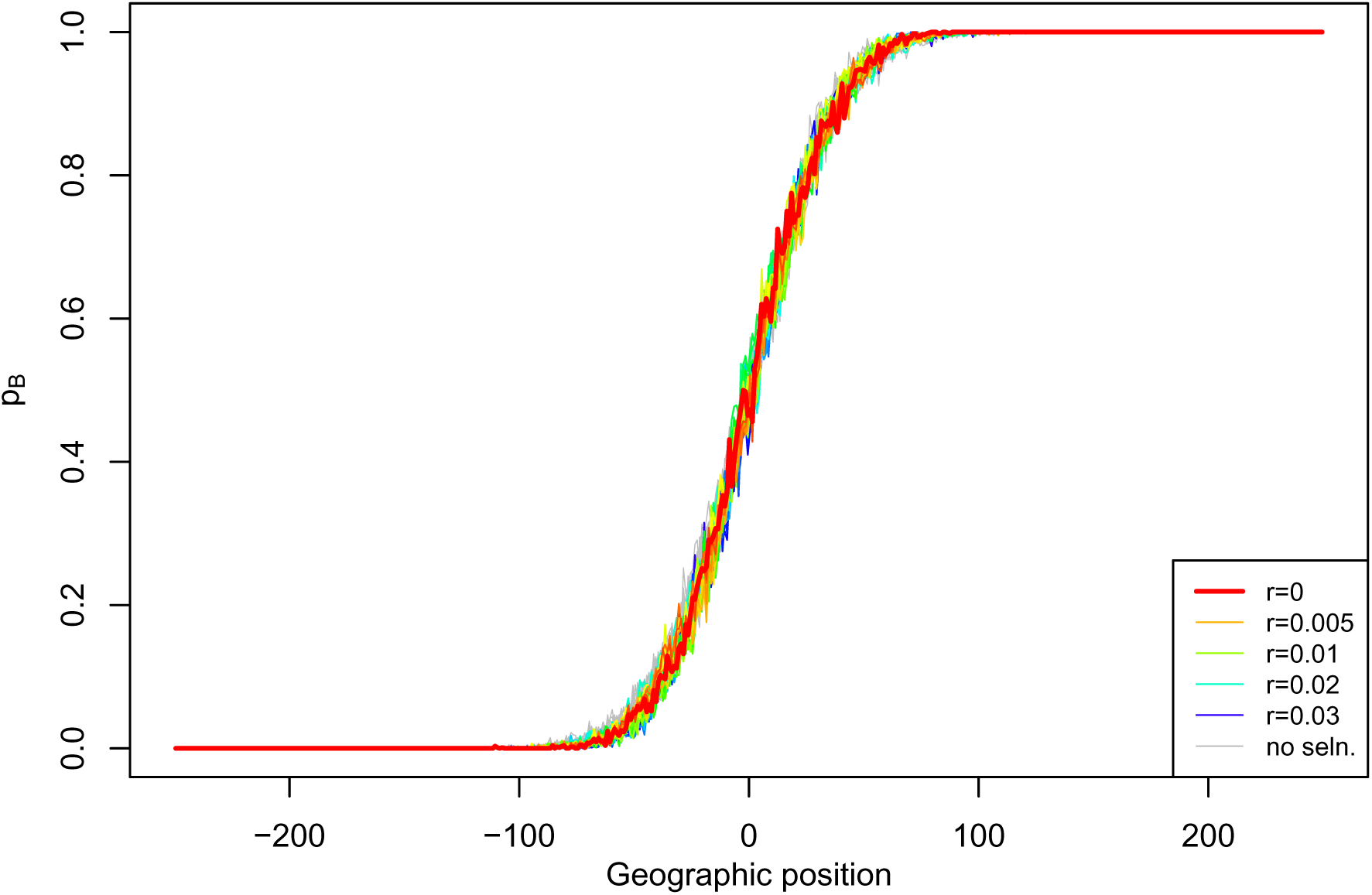
Frequency of ancestry *B*, across geography and at several physical positions on the genome, simulated for a hybrid zone *T* = 100 generations after secondary contact, and with *s* = 0.01. The simulated zone had 500 demes, each with a population size of 500 diploid individuals. Each line represents a locus some distance *r* away from the true target of selection, and *r* = 0 represents the locus that is under selection. Grey lines represent the same positions from a simulation with identical parameters except that *s* = 0.

**Figure S6:**
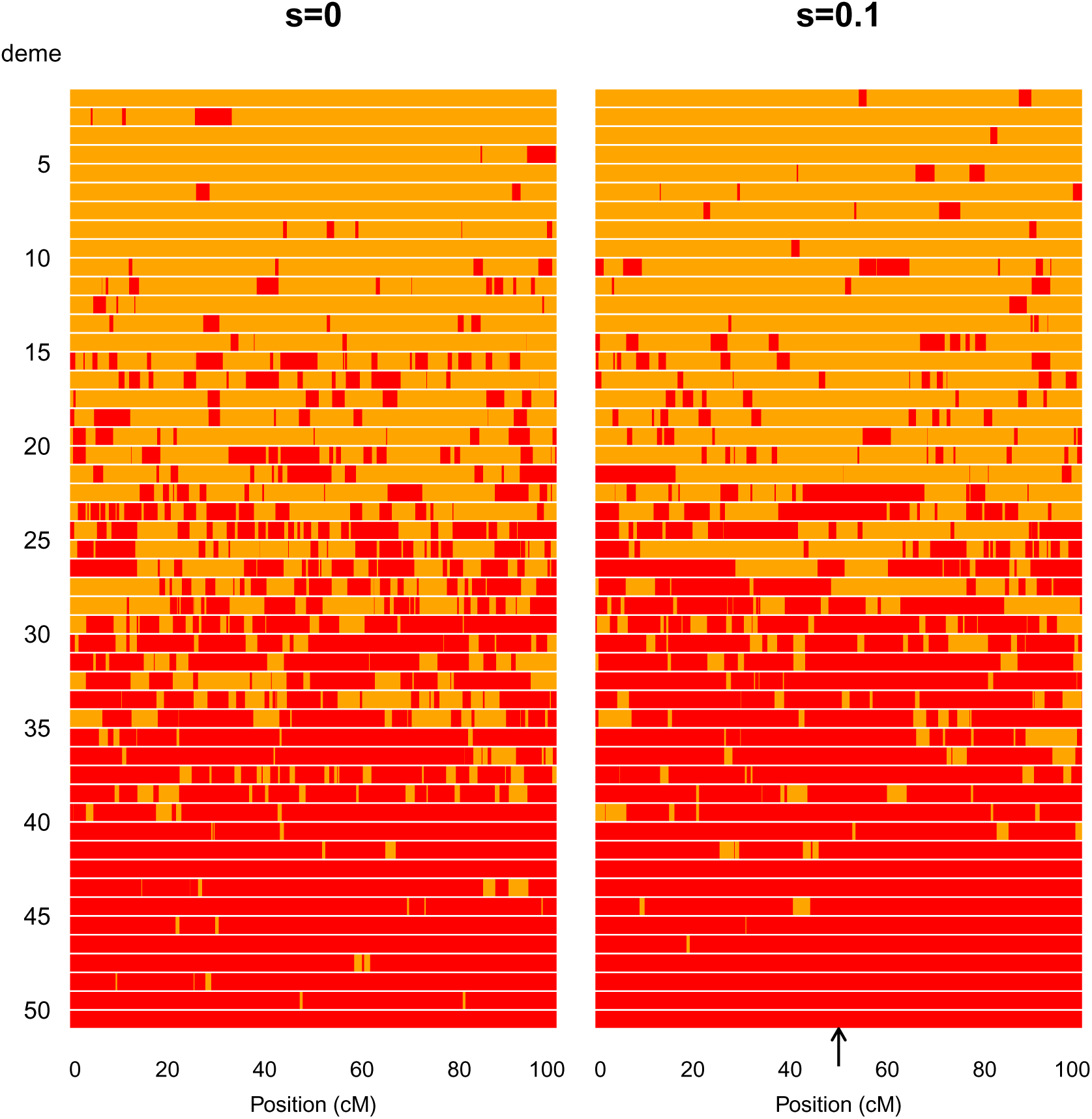
Randomly sampled chromosomes across a hybrid zone of age *T* = 100. Here we compare chromosomes of length 1M from a neutral zone to one that has a single under-dominant locus (*s* = 0.1) in the middle of the chromosome (indicated by black arrow). Red blocks along the chromosome denote ancestry *B*, and orange blocks are ancestry *A*.

**Figure S7:**
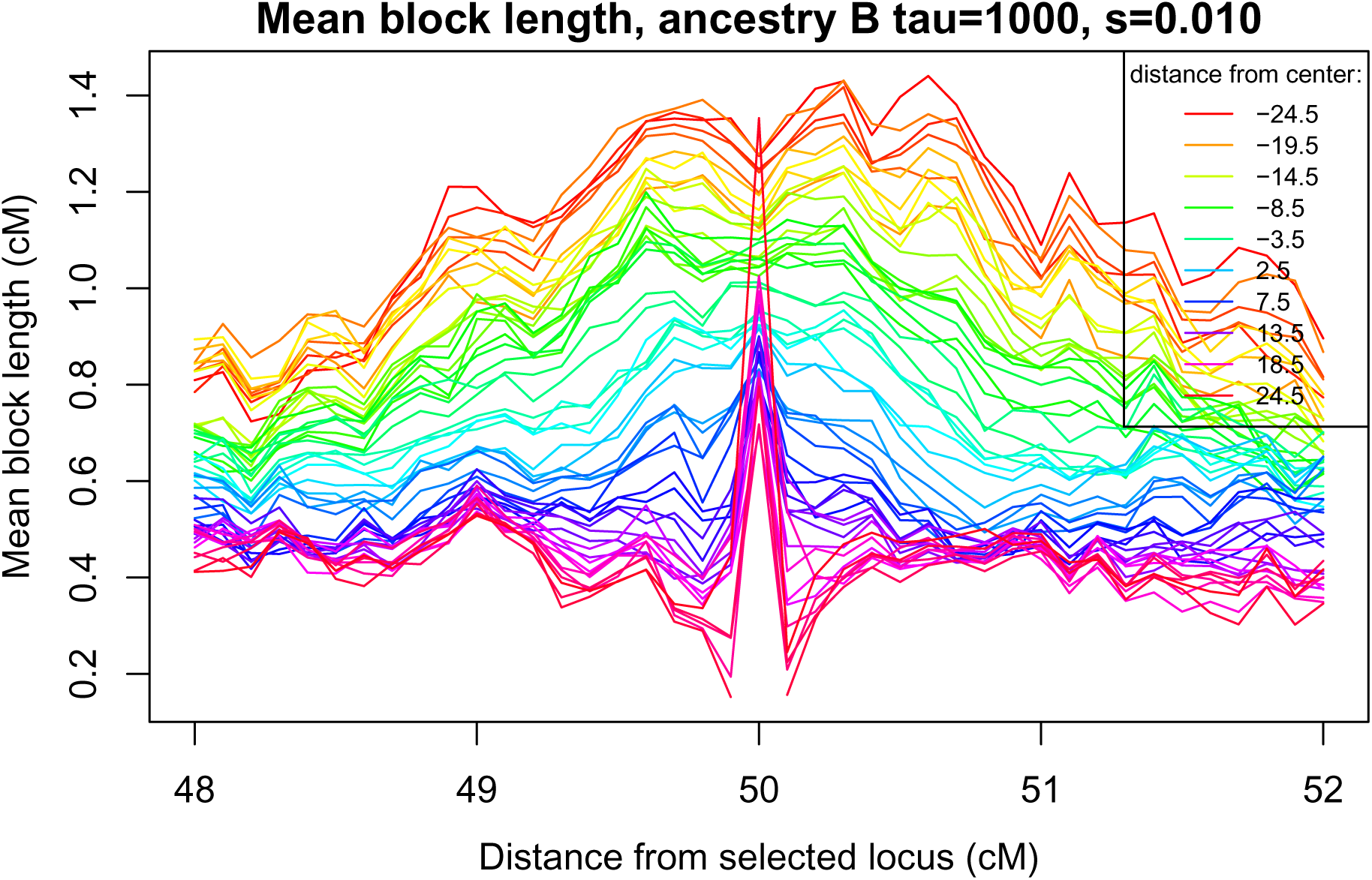
**Mean haplotype lengths of *B* haplotypes,** *l*_*B*_(*x*, *m*), across the genome (horizontal axis) and at different spatial locations (colored lines), from a simulation with 50 demes having 500 individuals each, *s* = 0.01, *σ* = 1, and after *T* = 1000 generations.

**Figure S8:**
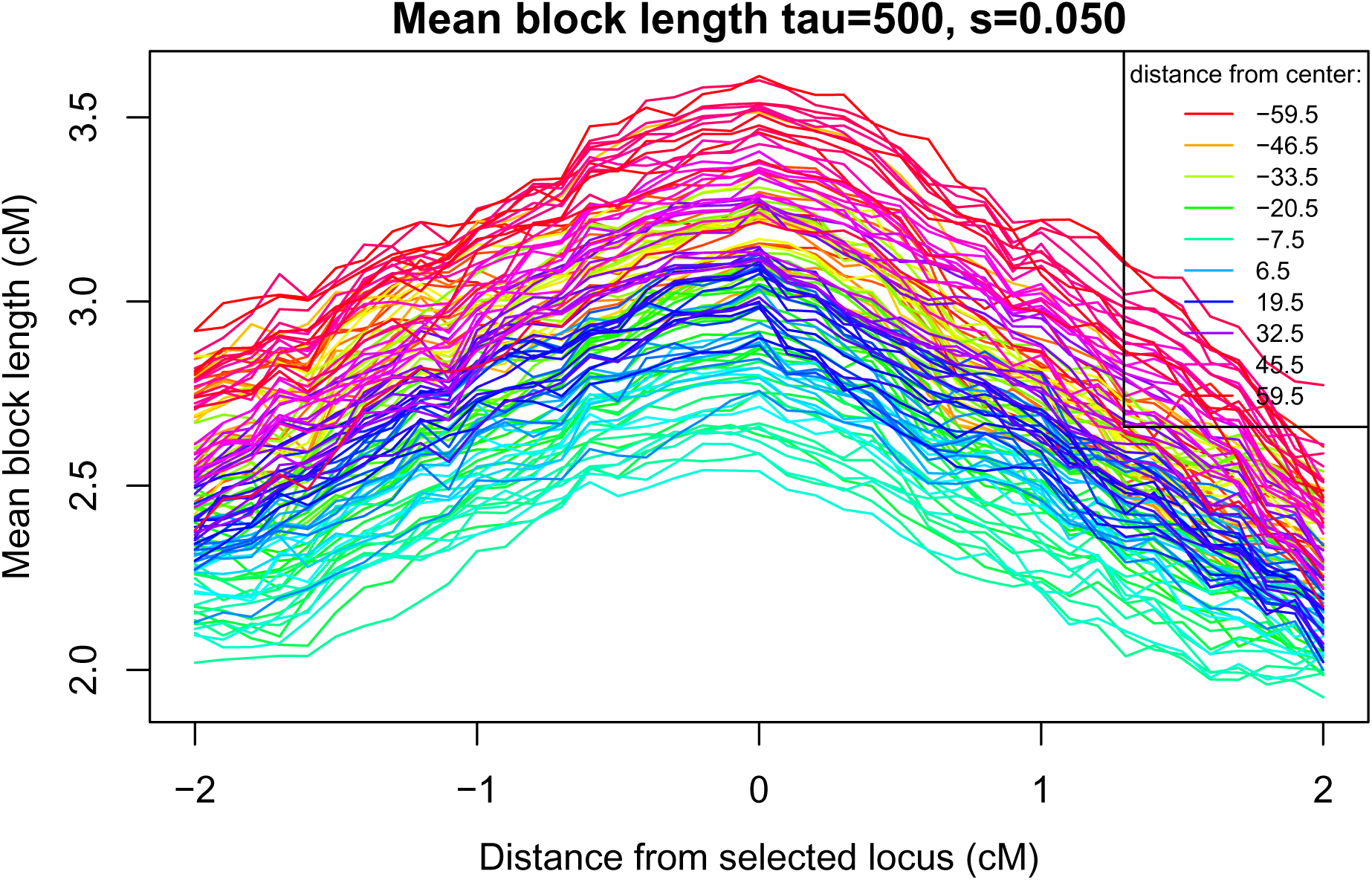
**Mean enclosing block length *l*(*m*, *x*),** across the genome (horizontal axis) and at different geographic positions (different colored lines). Results are from a simulation with 120 demes of 200 diploids each, selection *s* = .05, dispersal *σ* = 3, and after *T* = 500 generations.

**Figure S9:**
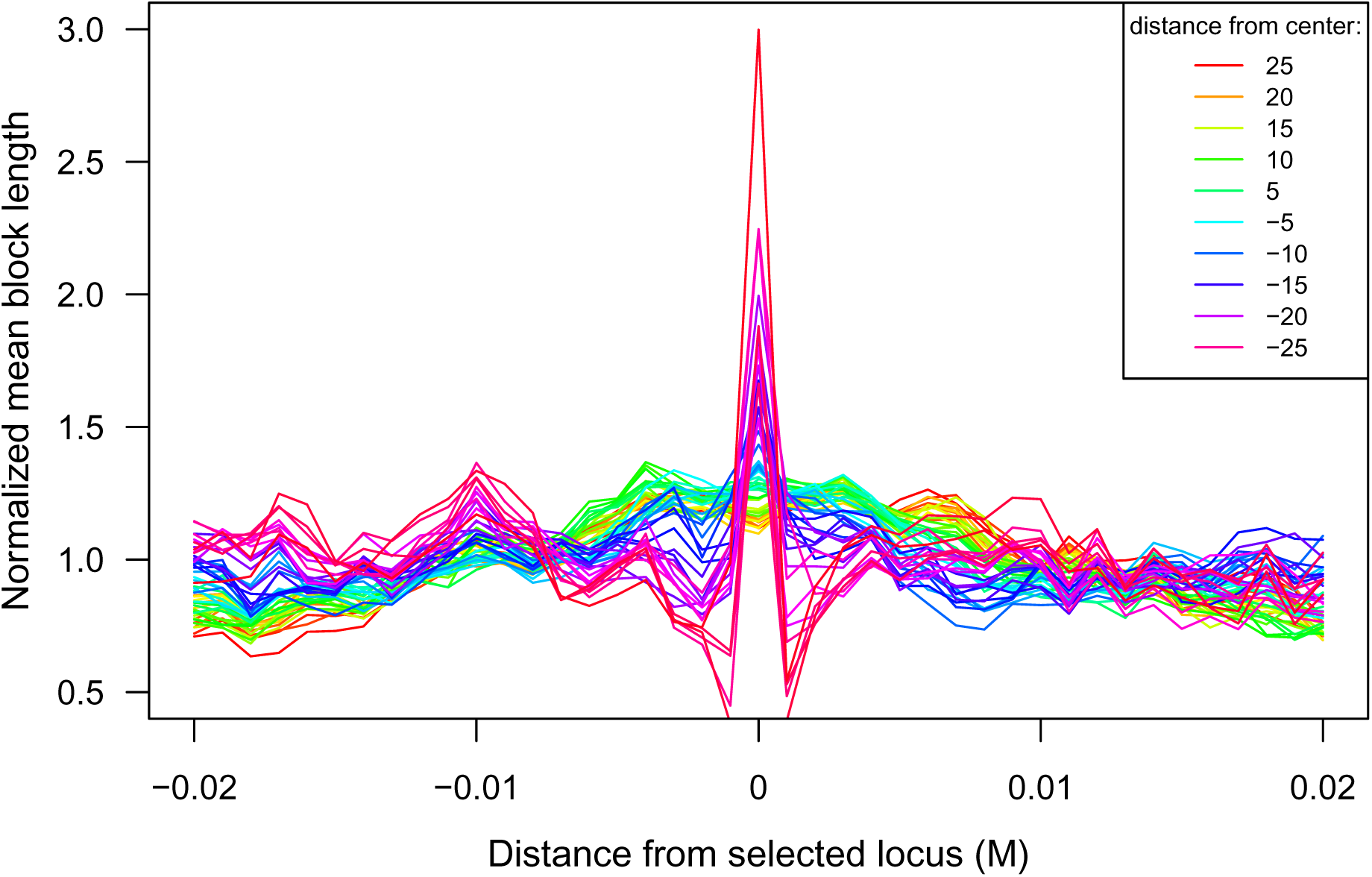
**Normalized mean enclosing block length,** *l*̅_*B*_(*m*, *x*), after *T* = 1000 generations, against position relative to the selected locus (horizontal axis) located in the center of a 1M chromosome. Each line shows the mean block length at that spatial and genomic position divided by the mean over the chromosome at that location; the simulation was run with *s* = 0.01 and *σ* = 1, 50 demes, each containing 500 diploid individuals.

**Figure S10:**
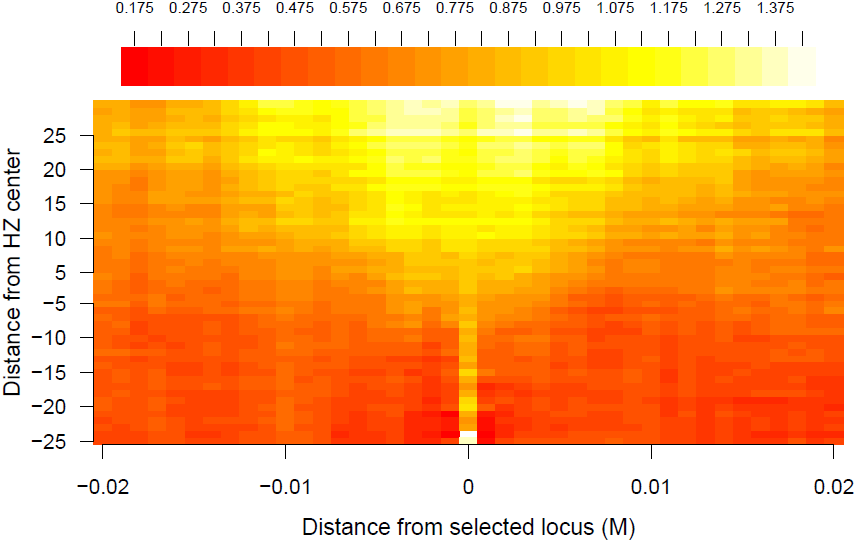
Heatmap of mean block length *l*_*B*_(*m*) along a simulated chromosome under *T* = 1000, *s* = 0.01 and *σ* = 1, 50 demes, each containing 500 diploid individuals.

**Figure S11:**
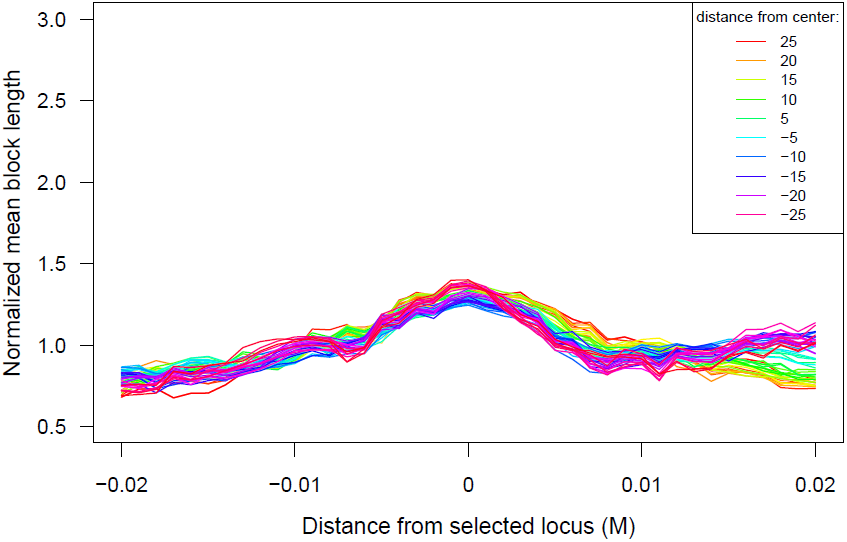
Mean block length *l*(*m*), without conditioning on ancestry, surrounding a given position along the genome with a single underdominant site with parameters as for Fig S9. (*s* = 0.01, *T* = 1000, *σ* = 1).

**Figure S12:**
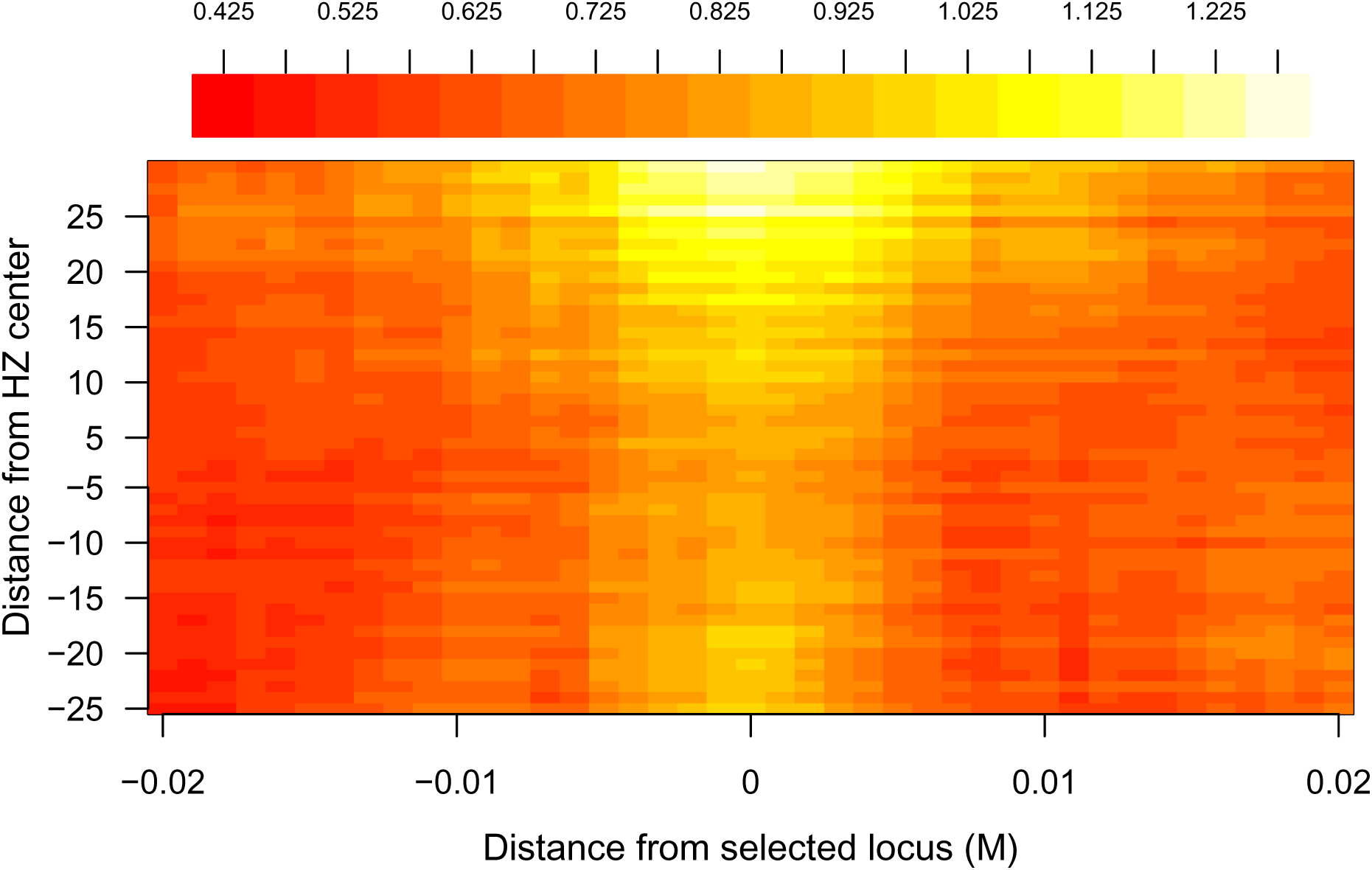
Heatmap of mean block length *l*(*m*), without conditioning on ancestry, along a simulated chromosome with a single underdominant site, with parameters as for Fig S9 (*s* = 0.01, *T* = 1000, *σ* = 1)

**Figure S13:**
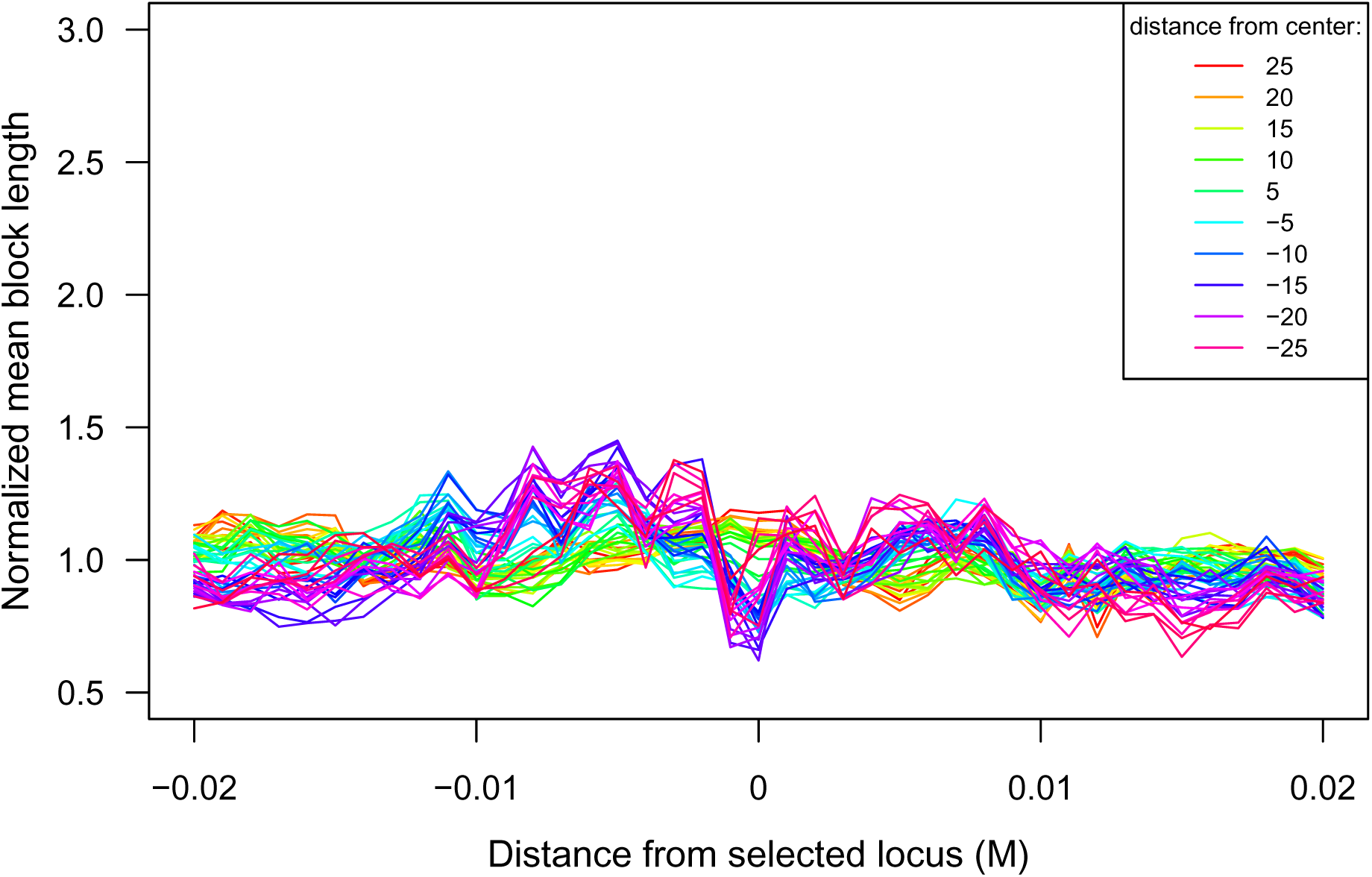
Mean block length of *l*_*A*_(*m*±) across chromosome with single under dominant site, conditioning on ancestry *B* at the selected locus, with parameters as for Fig S9 (*s* = 0.01, *T* = 1000, *σ* = 1)

**Figure S14:**
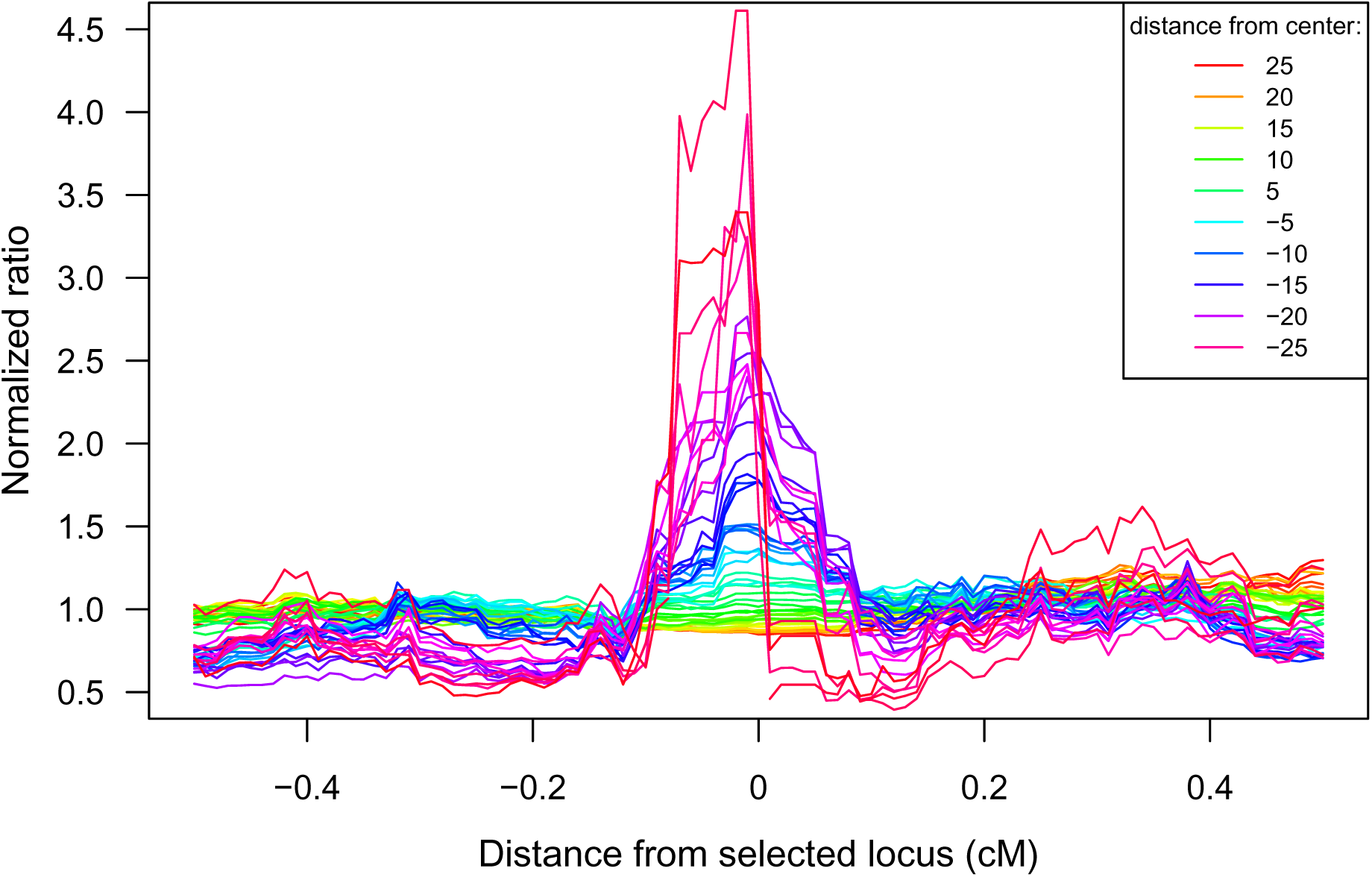
The normalized statistic *C*̅(*m*, *x*): the ratio 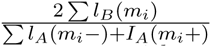 of mean block length and mean adjacent block lengths across a simulated chromosome with a single underdominant site and conditioning on ancestry *B* at the selected site (*s* = 0.01, *T* = 1000). Each line represents a deme and is normalized by mean block length across the chromosome in the deme. This simulation corresponds to that represented in Figure 5; note the difference in y-axis scale.

**Figure S15:**
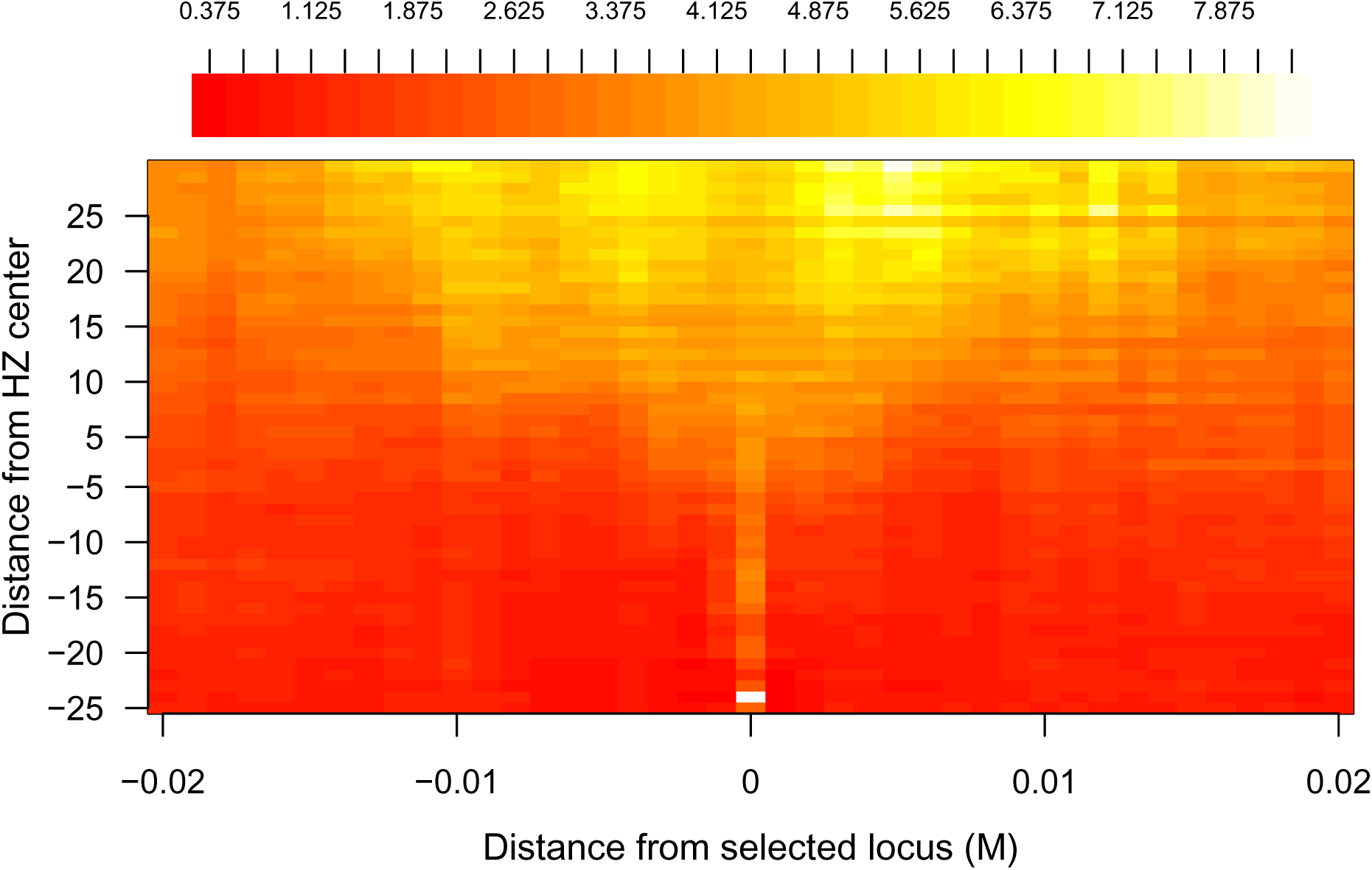
Heatmap of 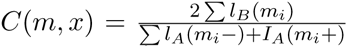 across a simulated chromosome with a single underdominant site and conditioning on ancestry *B* at the selected site and parameters as for Fig. S9 (*s* = 0.01, *T* = 1000, *σ* = 1).

**Figure S16:**
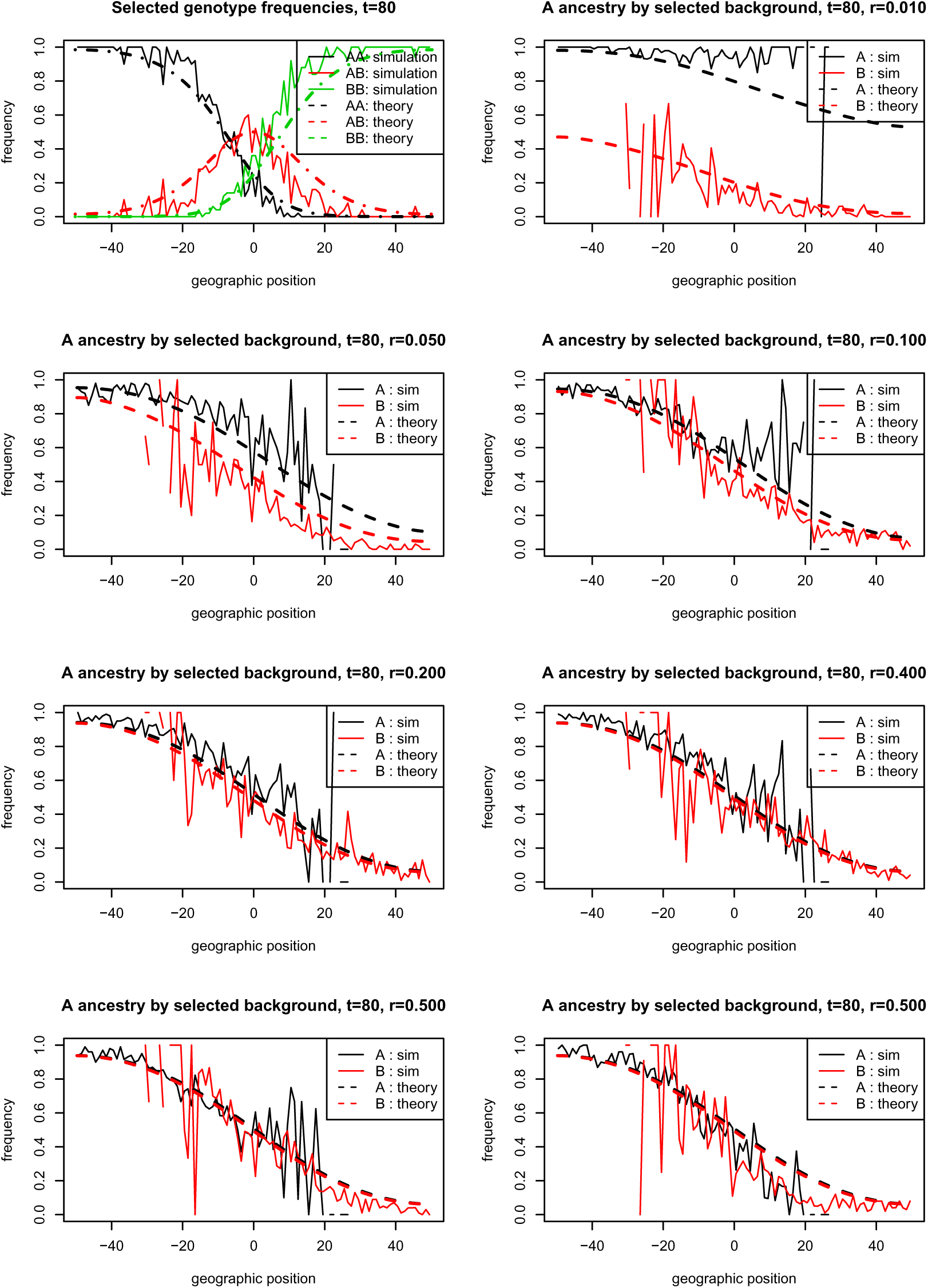
Conditional frequencies of ancestry *A* at *τ* = 80, comparing simulation and theory, with *σ* = 3 deme spacings, *s* = 0.05, and 50 individuals per deme. The top left figure shows observed and expected genotype frequencies for the two homozygotes and the heterozygote at the selected locus; expected genotype counts were obtained assuming random mating, and by solving equation (1) numerically. The remaining figures show observed and expected frequencies of *A* ancestry, separately conditioned on the identity of the linked allele at the selected site. Observed frequencies become much noisier where the linked allele becomes rarer.

**Figure S17:**
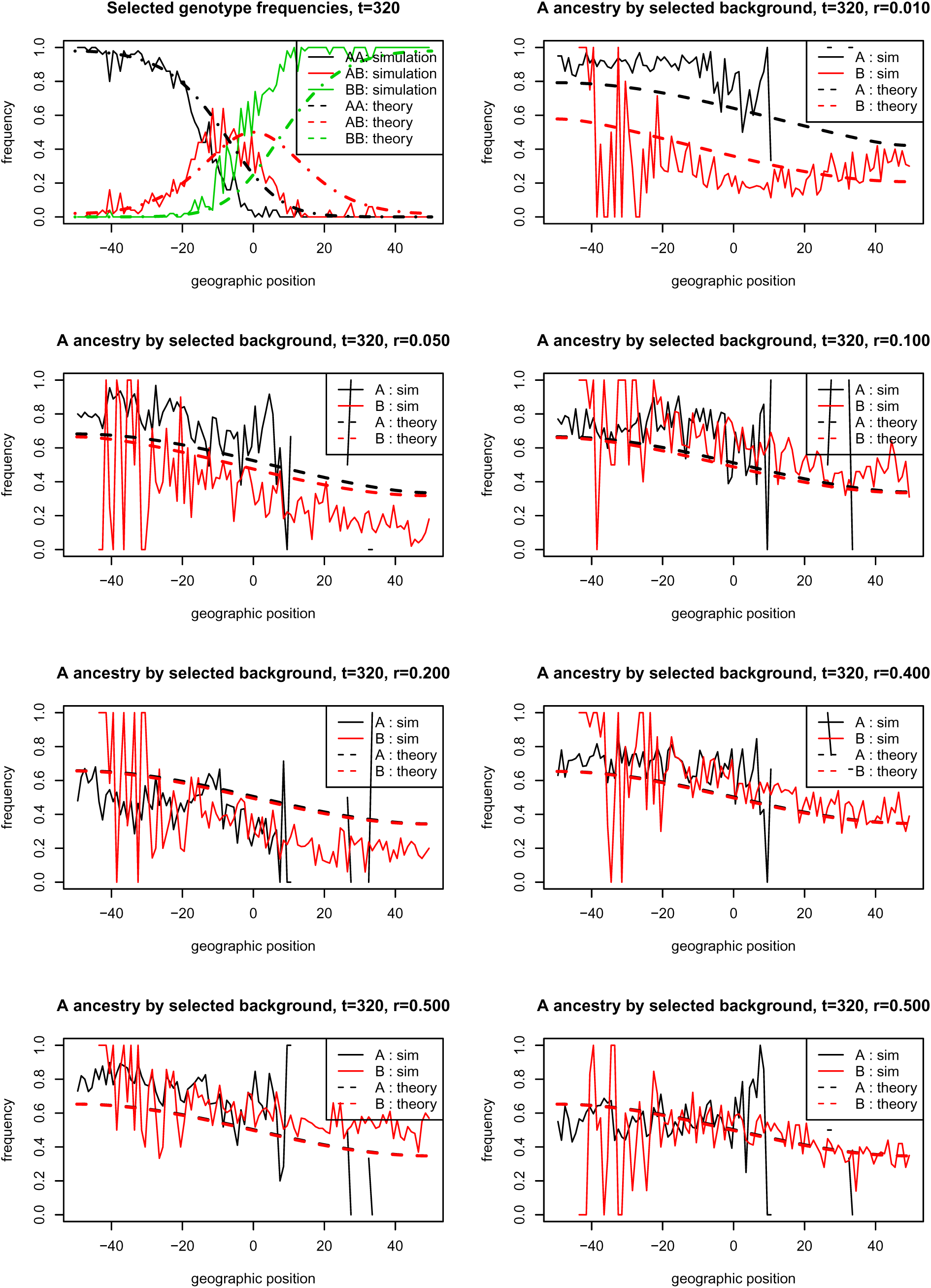
**Conditional frequencies of ancestry *A* at *τ* = 320,** as in figure S16. Deviations are larger than at *τ* = 80, due to genetic drift.

**Figure S18:**
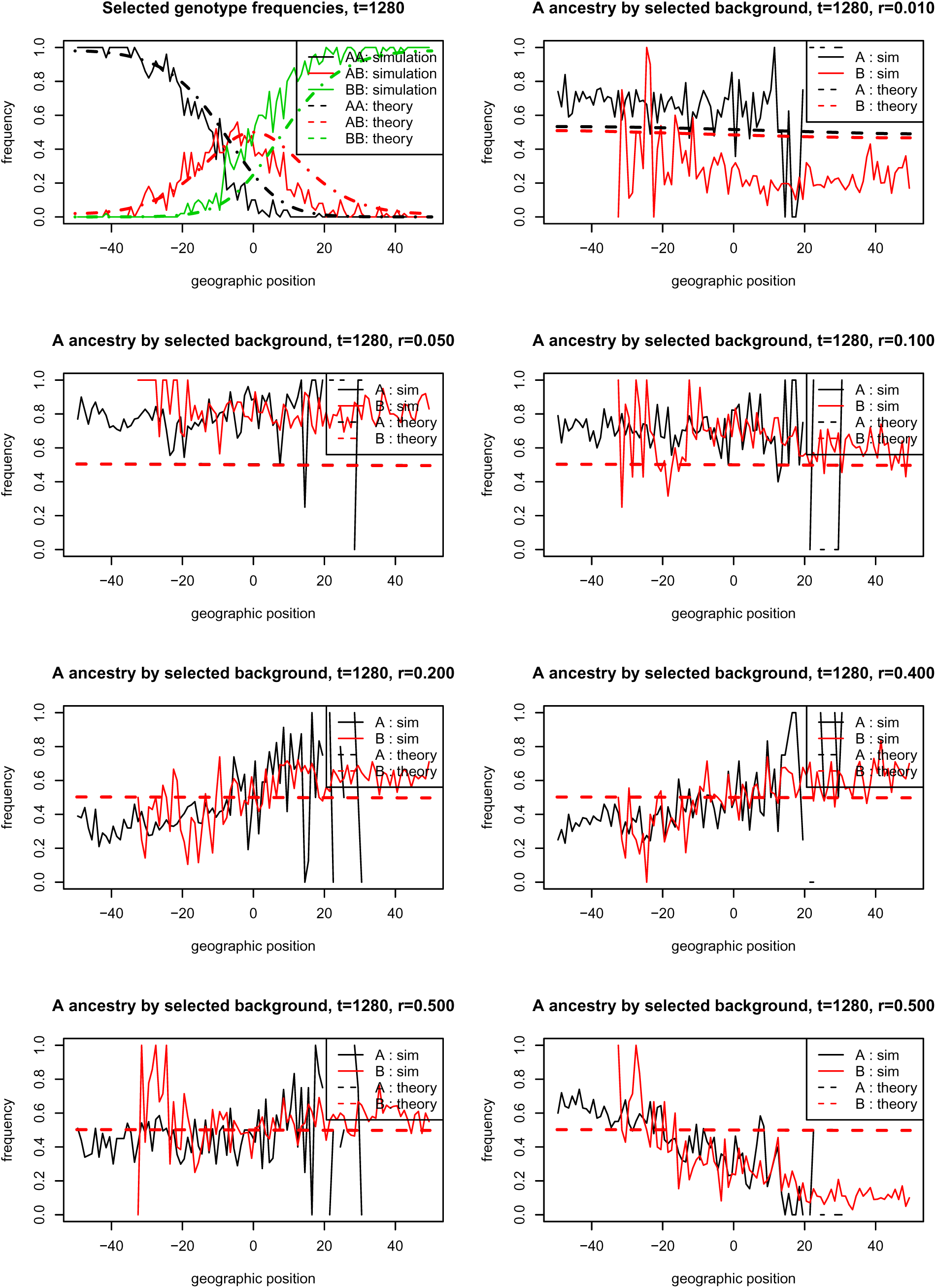
**Conditional frequencies of ancestry *A* at *τ* = 1280, as in figure S16.** Deviations are larger still than at τ = 320, due to genetic drift.

**Figure S19:**
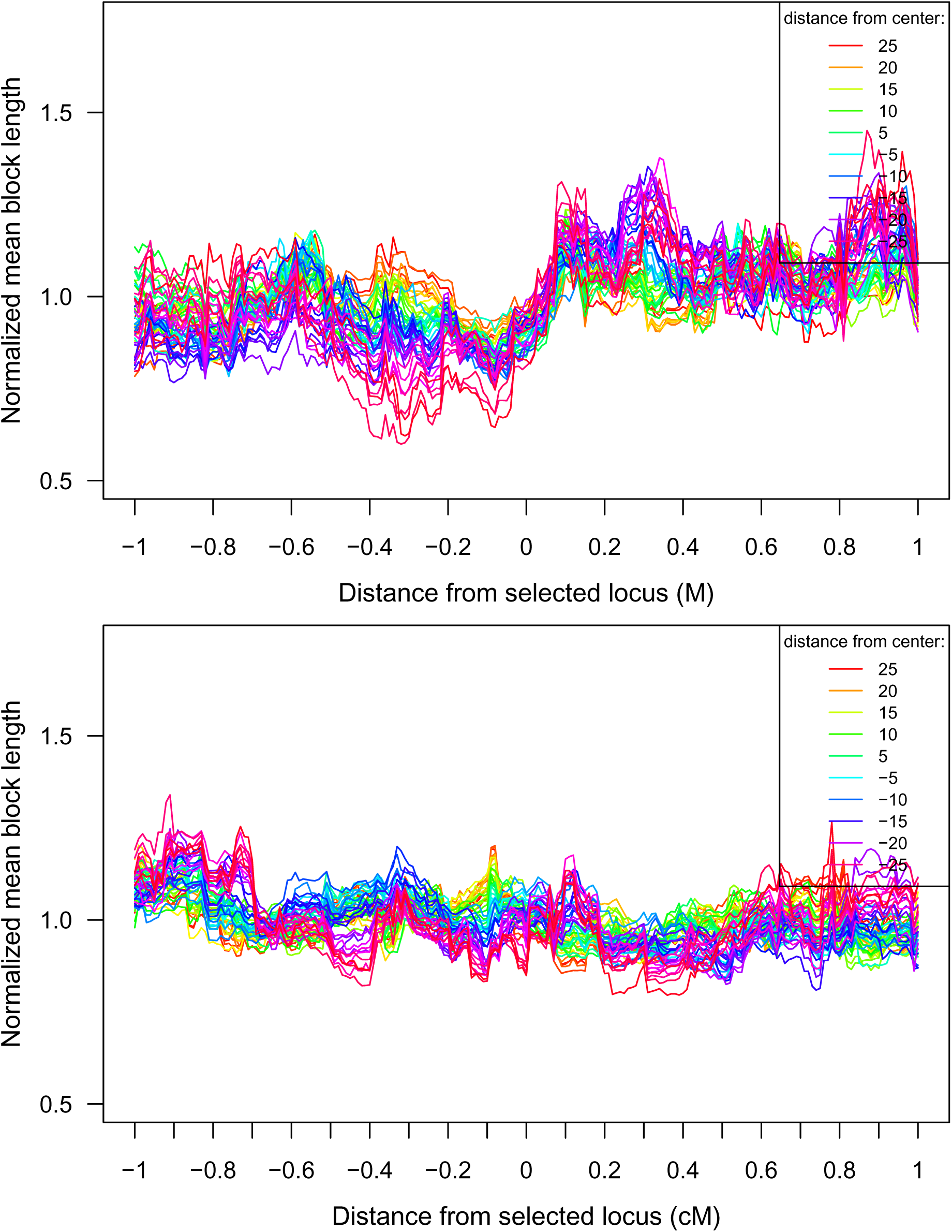
Normalized ancestry B block lengths (top) and all ancestry block lengths (bottom) along chromosome under *s* = 0.001 and other parameters as for Fig. S9 (*τ* = 1000, *σ* = 1).

